# Quantifying cooperative multisite binding in the hub protein LC8 through Bayesian inference

**DOI:** 10.1101/2022.06.29.498022

**Authors:** Aidan B Estelle, August George, Elisar J Barbar, Daniel M Zuckerman

**Affiliations:** Department of Biochemistry and Biophysics, Oregon State University, Corvallis, Oregon 97331, United States; Department of Biomedical Engineering, School of Medicine, Oregon Health and Science University, Portland, Oregon 97239, United states

## Abstract

Multistep protein-protein interactions underlie most biological processes, but their characterization through methods such as isothermal titration calorimetry (ITC) is largely confined to simple models that provide little information on the intermediate, individual steps. In this study, we primarily examine the essential hub protein LC8, a small dimer that binds disordered regions of 100+ client proteins in two symmetrical grooves at the dimer interface. Mechanistic details of LC8 binding have remained elusive, hampered in part by ITC data analyses employing simple models that treat bivalent binding as a single event with a single binding affinity. We build on existing Bayesian ITC approaches to quantify thermodynamic parameters for multi-site binding interactions impacted by significant uncertainty in protein concentration. Using a two-site binding model, we model LC8 binding and identify positive cooperativity with high confidence for multiple client peptides. Application of an identical model to two-site binding between the coiled-coil dimer NudE and the intermediate chain of dynein reveals little evidence of cooperativity, in contrast to LC8. We propose that cooperativity in the LC8 system drives the formation of saturated 2:2 bound states, which play a functional role in many LC8 complexes. In addition to these system-specific findings, our work advances general ITC analysis in two ways. First, we describe a previously unrecognized mathematical ambiguity in concentrations in standard binding models and clarify how it impacts the precision with which binding parameters can be determined in cases of high uncertainty in analyte concentrations. Second, building on observations in the LC8 system, we develop a system-agnostic heat map of practical parameter identifiability calculated from synthetic data which demonstrates that certain binding parameters intrinsically inflate parameter uncertainty in ITC analysis, independent of experimental uncertainties.

**Author Summary:** Multi-site protein-protein interactions govern many protein functions throughout the cell. Precise determination of thermodynamic constants of multi-site binding is a significant biophysical challenge, however. The application of complex models to multi-step interactions is difficult and hampered further by complications arising from uncertainty in analyte concentrations. To address these issues, we utilize Bayesian statistical techniques which calculate the ‘likelihood’ of parameters giving rise to experimental observations to build probability density distributions for thermodynamic parameters of binding. To demonstrate the method and improve our understanding how the hub protein LC8 promotes dimerization of its 100+ binding partners, we test the pipeline on several of these partners and demonstrate that LC8 can bind clients cooperatively, driving interactions towards a ‘fully bound’ functional state. We additionally examine an interaction between the dimer NudE and the intermediate chain of dynein, which does not appear to bind with cooperativity. Our work provides a solid foundation for future analysis of more complicated binding interactions, including oligomeric complexes formed between LC8 and clients with multiple LC8-binding sites.

## Introduction

Intracellular processes frequently depend on complex, multistep interactions between proteins or between proteins and small-molecule ligands(1–3). The hub protein LC8 provides an extreme example of binding complexity, accommodating over 100 client proteins via two symmetrical binding grooves(4,5) – often binding in multivalent fashion with a range of stoichiometries(6–10). LC8 is found throughout the eukaryotic cell and is involved in a host of cell functions, with client proteins including transcription factors(7,9), tumor suppressors and oncogenes(11,12), viral proteins(13–15), and cytoskeletal proteins(6,16).

Structurally, LC8 forms a small 20 kDa homodimer (Fig. 1a), with two identical binding grooves formed at the dimer interface(4,5). These binding sites induce a beta-strand structure in a well-characterized linear motif anchored by a TQT amino acid sequence within disordered regions of client proteins(6,9). Despite extensive study(9,16,17), the mechanistic thermodynamics of LC8 binding are still not fully understood, due to the difficulty of deconvoluting a multiplicity of microscopic states in its binding processes.

**Figure 1:**
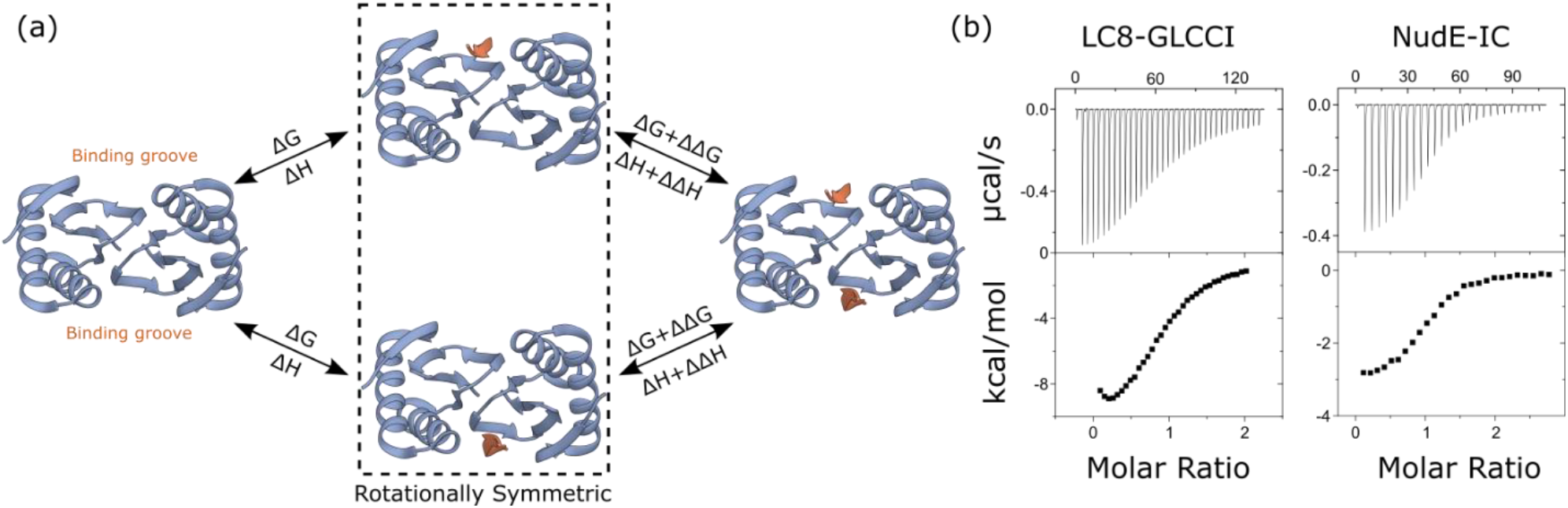
LC8 binds clients through a two-site mechanism. (a) Diagram of LC8-client binding, showing a structure of apo LC8 on the left, and a fully bound structure (PDB 3E2B) on the right. Intermediates are boxed to indicate they are symmetric and indistinguishable species. (b) Example isotherms for binding between dimeric LC8 and client peptide taken from GLCCI (left) and between the coiled-coil protein NudE and its client, the intermediate chain (IC) of dynein (right).

While published isothermal titration calorimetry (ITC) experiments fit LC8-client interactions to a simple model(9,16,17), binding in fact occurs in two distinct steps (Fig. 1a). Evidence of this was first observed in nuclear magnetic resonance (NMR) titrations of peptides into LC8, where, for some clients, a partially bound intermediate state was observable. Populations for each state were fit to a two-site model, which suggested weak cooperativity in binding, although statistically rigorous investigation was not undertaken(18). Further evidence that LC8 binding cannot be explained by a simple model emerged from ITC experiments that exhibit non-sigmoidal behavior, where rather than forming a plateau, early injections dip in heat per injection, forming a U shape at the beginning of the titration (Fig. 1b)(9). Single-site models, such as the identical sites model in Origin software(19), return strictly sigmoidal binding curves and are unable to fit the U shape in such isotherms. Although previous investigation of LC8 binding partners(9) fitted isotherms to an identical sites model, that study did not probe details of binding thermodynamics. The observed non-sigmoidal behavior clearly raises the possibility that these isotherms may fit well to a two-site model of binding more representative of expectations for LC8’s two client sites(20).

The use of ITC to interrogate complex systems and multisite binding is challenging, as ITC data is of relatively low information, and individual isotherms often fit well to varied model parameters(20,21). Despite this, ITC experiments can measure cooperativity(22,23), entropy-enthalpy compensation(17,24), changes in protonation state(25,26), and competition between multiple ligands(27,28). In general, these studies rely on fitting data globally to a model that includes several isotherms collected at varied conditions to reduce ambiguity of fit parameters(20,21), or a ‘divide and conquer’ type approach, where subsections of a complex binding network can be isolated and examined (16,22).

Concentration uncertainty is a critical concern in analysis of ITC data. In principle, accurate determination of protein and ligand concentration is a prerequisite for obtaining reliable thermodynamic quantities, yet these values are challenging to obtain for many systems(29–33). The most common software package for fitting ITC data, built into the data analysis and fitting software Origin, and distributed with calorimeters, attempts to account for this uncertainty in several models through the stoichiometric parameter *n*, which can fit to non-integer values to correct for error in macromolecule concentrations(19,34). However, in addition to assuming independent sites, this implementation ignores uncertainty in concentration of the titrant in the syringe, which is treated as a fixed value. The popular and highly flexible fitting software SEDPHAT greatly improves on Origin’s capabilities, allowing for both explicit or implicit (i.e. an ‘inactive fraction’ correction) uncertainty corrections(21,35). As the authors note, however, allowing for variation in both analyte concentrations makes binding constants indeterminable within SEDPHAT due to correlative effects among model parameters.

Bayesian analysis offers a natural framework for incorporating uncertainty in concentration measurements in ITC analysis(32,36). In a Bayesian framework, thermodynamic parameter determination is guided by a mix of experimental data and ‘prior’ information, such as uncertainty ranges/models, that weights the overall ‘posterior’ probability of a given set of thermodynamic parameters. The posterior distribution of estimated binding parameters generated through Bayesian analysis is a complete description of the probability range of each model parameter – and correlations among parameters – based on the input data and priors. With a meaningful prior description of concentration uncertainty, there is reduced risk of underestimating uncertainty in thermodynamic binding parameters. The Bayesian framework accounts for the full-dimensional likelihood of parameter space by construction, in contrast to maximum-likelihood approaches to uncertainty quantification in multi-parameter systems which approximate the likelihood function based on optimal parameters(21,37).

We build on earlier applications of Bayesian inference to ITC. Nguyen et al. (2018)(32) studied 1:1 binding using a Bayesian statistical framework accounting for concentration uncertainty and performed sensitivity analysis on concentration priors. For a two-site binding model, Duvvuri et al. (2018)(38) demonstrated that a Bayesian method can accurately and precisely determine two separate affinities when applied as a global model to several isotherms, but the work assumes no uncertainty in measured concentrations(38), raising the possibility that parameter uncertainty is underestimated(21,32). Cardoso et al. (2020)(36) used a simplified 4-site binding model with a single common binding enthalpy for a set of isotherms to determine 3 of 4 distinct affinities between protein and ligand, with the fourth being uncertain across a range of several orders of magnitude. Although Cardoso et al. (2020)(36) include concentrations as model parameters, they greatly narrow concentration priors using a preliminary ‘calibration’ assuming identical sites. We note that such a model is not appropriate for complex systems, particularly in cases where the identical-sites model does not fit well to the isotherm shape. A sensitivity analysis regarding concentration uncertainty was not performed in either multisite study, and neither work probed the information content of single isotherms for multisite systems.

Here, we report a Bayesian analysis of two-site systems with proper accounting of concentration effects critical for reliable analysis. We show that LC8-client interactions unambiguously exhibit positive cooperativity, driving binding towards a fully bound state. In contrast, symmetric two-site binding between the coiled coil domain of the dynein cargo adaptor NudE and the intermediate chain (IC) of dynein(39) shows no significant evidence for cooperativity.

We also provide methodological advances. First, we derive simple mathematical relations that govern the influence of concentration uncertainties on different binding parameters, providing a fundamental basis for the previously noted strong sensitivity of enthalpies – but not free energies – to concentration uncertainty(21). Second, by using synthetic models, we systematically characterize the causes of binding-parameter uncertainties in two ways: we demonstrate that substantial uncertainty can result from the binding parameters themselves, e.g., strong vs. weak binding; and we also determine the effects of different prior functional forms and uncertainty ranges in a multisite context, extending the work of Nguyen et al. (2018)(32). Finally, we outline best practices for determining model parameters and uncertainties in a multisite Bayesian framework.

## Results

### A mathematical “degeneracy” in thermodynamic parameters impacts analysis at any stoichiometry

We first present a simple mathematical analysis that explains previously reported correlation effects among titrant and titrand concentrations(21), and which significantly impacts the overall analysis of ITC data. Specifically, when the concentrations are uncertain, as is common in analysis of ITC data(21,32), we show below that only the *ratio* of titrant:titrand concentrations can be estimated, rather than the individual values, and this ambiguity propagates to all thermodynamic parameters. Hence, there is a “degeneracy” in that multiple solutions (sets of concentration values and thermodynamic parameters) will equally describe even idealized ITC data lacking experimental noise (Fig. 2).

**Figure 2:**
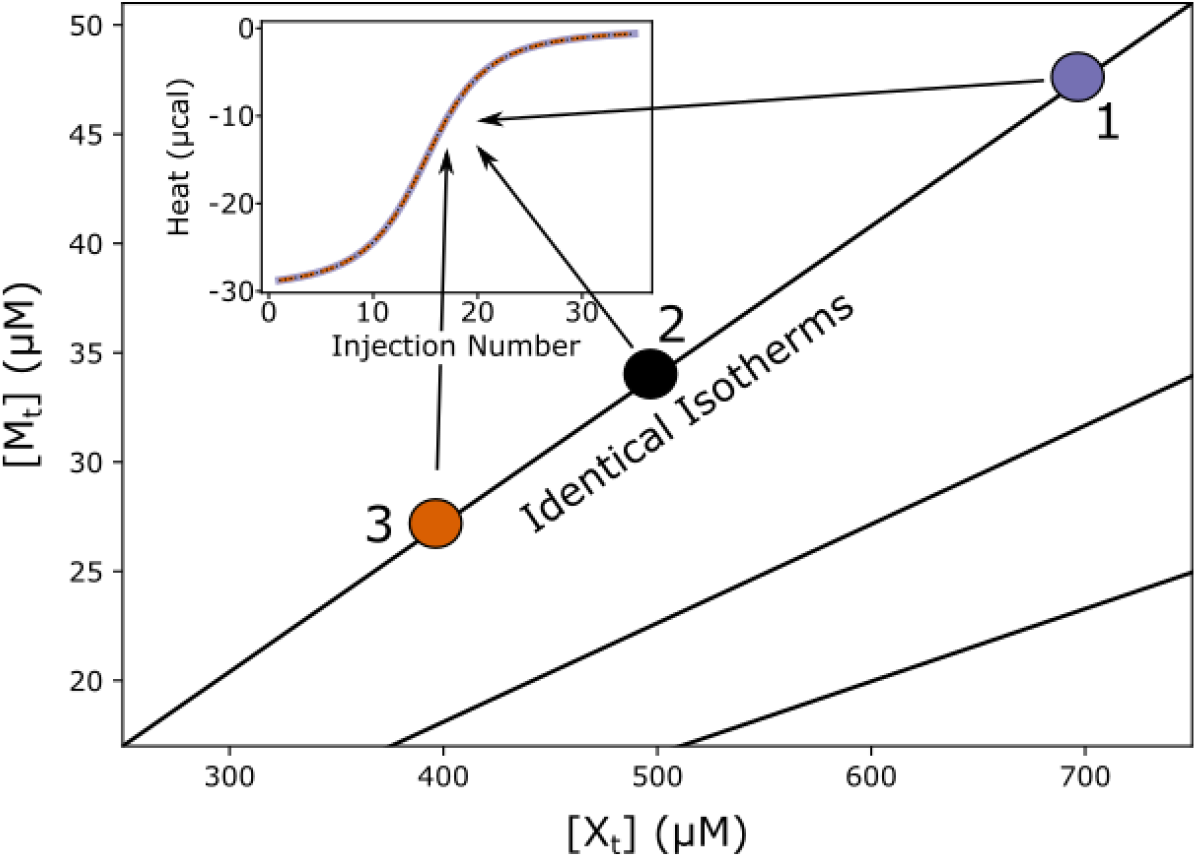
Exact degeneracy in binding isotherms. Based on the scaling relations of Eq (2), for any set of ligand and total macromolecule concentrations (X_t_, M_t_), there are infinitely many alternative concentrations (e.g., filled circles) on a diagonal line in the ([X_t_], [M_t_]) plane which yield exactly equivalent isotherms (inset, isotherms for points 1, 2, and 3 are drawn in distinct colors but overlay exactly) for a fixed set of thermodynamic parameters. For any given point in parameter space, equivalent degenerate lines can be drawn in a radial manner (e.g. the two additional black lines), passing through the point and the origin. The plotted synthetic isotherms are for 1:1 binding, but analogous degeneracy also holds for multivalent binding - see text. Note that no fixed relationship among concentrations is assumed during Bayesian inference.

We first describe the degeneracy for standard 1:1 binding between a macromolecule M and ligand X, following the scheme

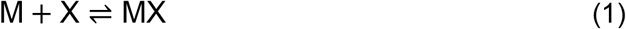

The heat, Q, of a 1:1 binding system at any titration point can be described using the standard quadratic binding equation used in the identical sites model(19,40):

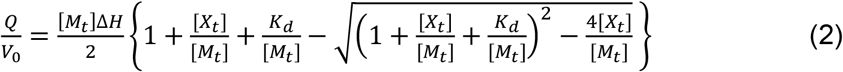

where [M_t_] and [X_t_] are the concentrations of macromolecule and ligand (i.e., cell component and syringe component) respectively, while K_d_ and ΔH are the binding affinity and enthalpy. The degeneracy is demonstrated by introducing a linear scaling of all parameters by an arbitrary number denoted *α*. Specifically, we apply the following transformations:

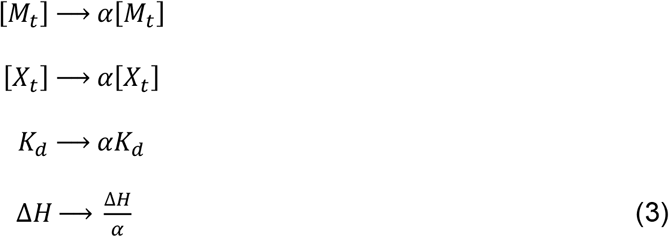

Applying this set of transformations, we can rewrite the binding equation:

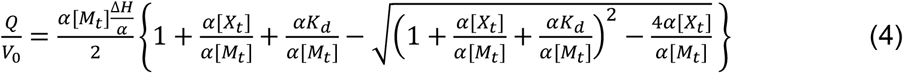

Regardless of the value of the factor α, all introduced factors cancel leaving Q unchanged.

We emphasize that the factor α is used here as a mathematical tool to analyze the governing equations, but α is *not* a parameter in our Bayesian inference pipeline, and we do *not* assume analyte concentrations bear any fixed relation to one another in the inference pipeline. In the Bayesian inference process, parameters can take any values within the ranges allowed by the priors.

Nearly identical formal considerations apply in the two-site binding model of primary interest here. As detailed in the Methods, the value of Q is unchanged when both concentrations and both K_d_ values are multiplied by α and both ΔH values are divided by α. The underlying model is more complex as it requires solving a system of nonlinear equations (see Methods for details), but the result is that α is propagated through the nonlinear equation solutions, and once again cancels in the calculation of Q, leaving the heat value unchanged.

To facilitate analysis and discussion of cooperativity below, we parameterize two-site our model using ΔG, ΔΔG, ΔH and ΔΔH. Values are ‘microscopic’ terms corresponding to a single site-resolved binding step. The ΔΔG and ΔΔH value correspond to the differences between the first and second microscopic binding steps. Thus K_d1_ = e^ΔG/RT^, K_d2_ = e^(ΔG+ΔΔG)/RT^, and ΔH_2_ - ΔH_1_ = ΔΔH. The energy-like formulation allows for easy assessment of cooperativity (ΔΔG will be zero in the absence of cooperativity and positive or negative for negative or positive cooperativity, respectively), and ΔΔH is the change in enthalpy between binding steps with analogous characterization.

The degeneracy and associated scaling relationships in Eq (3) provide important insight into assessment of thermodynamic parameters inferred from ITC data. We see directly that binding enthalpy changes proportionately to concentrations of titrant and titrand. That is, a given percent error in an assumed concentration of either ligand (characterized by alpha) translates to the same scale of error in ΔH. On the other hand, the binding free energy ΔG, is less sensitive to concentration errors, due to scaling with ln(α), rather than directly multiplied by α. We note again that α is used only in the formal analysis here and not in our Bayesian inference process where parameters are not assumed to have any fixed relation with one another.

The scaling relationships of Eq. (3) also presage a significant issue in Bayesian inference, namely, sensitivity to the choice of priors. Within the set of degenerate solutions (diagonal lines of concentration pairs in Fig. 2), the Bayesian ‘likelihood’ probability – which describes how well a parameter set fits the data in the absence of prior information – will be constant, as solutions are mathematically identical. Thus, within any degenerate set, the assumed prior distributions for concentrations, will determine the overall posterior distributions (see Methods). Because the posterior distributions ultimately determine the uncertainty ranges, this is a key point.

Below, we continue to examine the ramifications of the concentration degeneracy, demonstrating concretely that enthalpy is more impacted by uncertainty in concentrations than free energy. We also examine the influence of priors on parameter distributions and discuss parameter distributions determined from isotherms in cases of high concentration uncertainty.

### Validation of Bayesian inference pipeline with synthetic data

To test our Bayesian pipeline (Methods), we generated ‘synthetic’ simulated isotherms using hand-chosen sets of thermodynamic parameters ΔG, ΔΔG, ΔH, ΔΔH (see Fig. 1) inserted in Methods Eq. 17 with added Gaussian noise. Following an exploration using synthetic data of how cooperativity impacts binding isotherms (e.g. Fig. 3a), we selected synthetic model parameters to mimic the isotherm shape seen in LC8-peptide binding examples. Specifically, slight positive cooperativity (ΔΔG = -1, ΔΔH = -1.5 kcal/mol) was best-suited to imitating real LC8-peptide isotherms, along with ΔG = -7 and ΔH = -10 kcal/mol. Synthetic noise is taken from a Gaussian distribution with a zero mean and standard deviation σ = 0.2 μcal. As shown in Fig. 3, we used our pipeline to sample posterior distributions for these isotherms. For concentrations, we chose uniform prior distributions of ±10% of the true value (which simply limits sampled concentration values to these ranges). The choice of 10% approximates what we view to be an attainable level of uncertainty for experimental protein concentrations.

**Figure 3:**
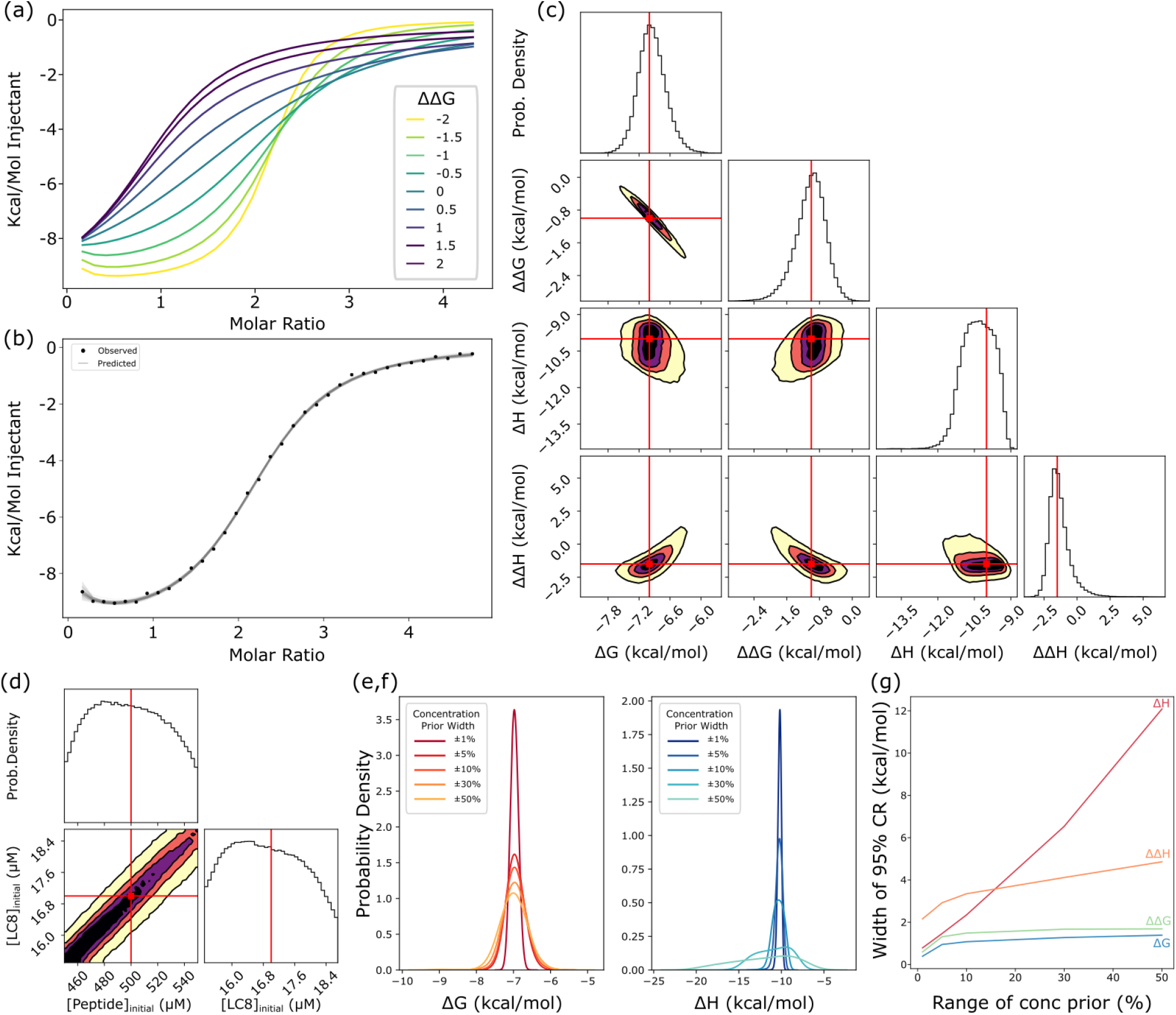
Analysis of two-site model using synthetic isotherms. (a) A set of synthetic isotherms for two-site binding with varied ΔΔG parameters demonstrating how cooperativity changes isotherm shape. Thermodynamic parameters are ΔG = -7, ΔH=-10, and ΔΔH=0. Concentrations are set at 17 and 500 μM for cell and syringe respectively, and injection volumes are 6 μL. (b) A synthetic isotherm (ΔG = -7, ΔH = -10, ΔΔG = -1, ΔΔH = -1.5 kcal/mol) with added gaussian noise (points) with 50 fitted isotherms (lines) generated through the Bayesian pipeline, i.e., sampled from the posterior. (c) One and two-dimensional marginal distributions for thermodynamic parameters, with contours in the two-dimensional plots set at 95 (yellow), 75 (orange), 50(purple) and 25%(black) confidence. Red lines and dots indicate true values for the synthetic isotherm. Marginal distributions, along with MCMC chains for all eight model parameters, including nuisance parameters can be found in Supplementary Figure S1. (d) Marginal distributions for concentration parameters, exhibiting characteristic diagonal shape (Fig. 2) with contours as in (c). Plots at the top of each column in panels c and d are one-dimensional probability density distributions. (e,f) One-dimensional distributions for ΔG (e) and ΔH (f) plotted for models with prior ranges for concentrations of 1, 5, 10, 30 and 50% of the stated concentration. (g) Width of the 95% Bayesian credibility region, akin to a confidence interval, for thermodynamic parameters as a function of the width of the concentration prior used in modeling, plotted from models with prior ranges for concentrations of ± 1, 5, 10, 30 and 50% of the stated concentration.

Under these representative conditions, inferred posterior distributions fell around the known model parameters, and model parameters equate to isotherms which closely matched the isotherm shape (Fig. 3b,c). The finite widths of the distributions are due both to synthetic experimental noise and correlative effects from the discussed model degeneracy. The posterior distribution for ΔG covers a range of ∼1 kcal/mol distributed around the true value of -7 kcal/mol. Examination of the distribution lets us define a ‘credibility region,’ that contains 95% of the distribution probability (i.e., from the 2.5 to 97.5%ile of the distribution), which is directly analogous to a confidence interval in frequentist terms. For ΔG, the 95% credibility region is -7.5 to -6.4 kcal/mol. Similarly, the 95% credibility region for ΔΔG covers a range of ∼1.5 kcal/mol, evenly distributed around -1 kcal/mol. ΔH and ΔΔH both have slightly wider credibility regions, with widths of 2.3 and 3.3 kcal/mol respectively, but both are distributed around the true values of -10 and - 1.5 kcal/mol respectively.

One benefit of Bayesian inference is the ability to examine multi-dimensional likelihood distributions to obtain correlations between model parameters without approximation. For example, in our two-dimensional distributions for the thermodynamic parameters, the ΔG and ΔΔG values are strongly negatively correlated (Fig. 3c), indicating a compensatory effect in the model, where increases in ΔG can be compensated by decreases in ΔΔG to arrive at similar solutions. Resultantly, the distribution for both ΔG and ΔΔG are broader than the ‘total’ free energy (i.e., 2*ΔG* + *ΔΔG*), evidence the overall energy of binding can be determined more precisely the energy of each step (S2 Fig.). Additionally, the mathematical degeneracy for concentrations described above can clearly be seen in these two-dimensional correlations: the two-dimensional marginal distribution for each concentration is a straight line of a width determined by noise covering the entire prior range (Fig. 3d). The scaling relationship of the model parameters outlined previously means that each point along this diagonal corresponds to a degenerate solution, i.e., each point has equivalent likelihood based on the data.

### Impact of concentration degeneracy on two-site thermodynamic parameters assessed via synthetic data

Bayesian inference enables determination of distributions for thermodynamic parameters even in cases of a concentration degeneracy. The net result, as will be seen, is a broadening of (posterior) parameter distributions based on multiple equally likely solutions, constrained by the priors used. Despite intrinsic limitations surrounding concentrations, the *ratio* of concentrations can be quantified with relatively high precision even when individual concentrations are highly uncertain. To quantify the impacts of the concentration degeneracy within a Bayesian inference pipeline, we examined a series of uniform prior distributions for concentrations, ranging from ±1% to ±50% for both concentrations. These priors were applied to a synthetic isotherm mimicking experimental parameters, as described in the pipeline validation above. The choice of concentration priors – which embody assumed or estimated experimental uncertainties – greatly impacts the predicted uncertainty of thermodynamic parameters. The distributions for ΔG and ΔH, not surprisingly, both widen as the prior range is increased (Fig. 3e,f). As anticipated by the degeneracy scaling relations of Eq (3), the width of the distributions for ΔH and ΔΔH increases roughly linearly with the concentration prior range, while the distributions for ΔG and ΔΔG increase initially at low concentration ranges then level off. This can be explained by the logarithmic relationship between the K_d_ (which is what scales with the degeneracy) and free energy. Functionally, high uncertainty in concentrations therefore only slightly increases uncertainty in binding free energy, while having a more significant impact on binding enthalpy.

The concentration degeneracy of the model limits the degree to which erroneously determined individual concentrations can be corrected. As discussed above, the fact that the Bayesian likelihood is uniform at any point along the degeneracy lines (Fig. 2) means that the data have little impact on the posterior distributions for *individual* concentrations, which instead takes the shape of the prior used. This can be seen in the model validation example (Fig. 3d), where the posterior distribution is approximately uniform, echoing the uniform prior.

The ratio of concentrations (‘macromolecule’ to ‘ligand’), on the other hand, is a determinable parameter, as the ratio does not change along the degenerate line. Posterior distributions are therefore limited to this ratio. For example, when we sample the posterior for the same isotherm, but use a normal (i.e. Gaussian) distribution for one concentration prior and a uniform distribution for the other, both posteriors take the shape of a normal distribution (S3 Fig.). This is a direct result of the degeneracy identified above. Supplementary Table S1 shows concentration ratio credibility regions for the experimental systems. Because of the nearly determinative relationship between the prior and posterior concentration distributions, we elected to use uniform priors for concentrations throughout this work to avoid undue influence on our results from model priors.

For completeness, we also examined 1:1 binding with synthetic data. Overall, the impact of the concentration degeneracy on model parameters is similar (S4 Fig.): binding enthalpy posterior distributions are wider than free energy distributions. In response to changes in concentration prior ranges, the posterior for ΔG is more impacted than in the two-site model, but the distribution remains much narrower than that of the enthalpy, as in the two-site model.

### Application to 2:2 LC8:IDP Systems

We applied the Bayesian analysis pipeline to a set of 7 experimental isotherms of binding between LC8 and client peptides, all of which bind in a 2:2 ratio. Note that the two LC8’s form a strong homodimer (Kd ∼ 60 nM)(41) and this initial homodimer formation is excluded from our analysis. Client peptides were chosen based on a K_d_ of less than 5 μM and deviation from the standard sigmoidal isotherm shape(9). As noted above, the user-supplied uncertainties for concentrations may impact uncertainty in other parameters. Following analysis with priors of ±10% and ±20% of the measured LC8 concentration as determined by absorbance at 280 nm, we have elected to focus on results at ±10% (Table 1), as moving to ±20% does not greatly alter the posterior distributions (S2 Table). The high degree of purity of LC8 (>95%) and high absorbance at 280 nm, due to the presence of 6 chromophores (1 Trp, 5 Tyr) allow for a high signal-to-noise ratio for the absorbance, reducing uncertainty in the measurement. Comparatively, because of the difficulty in accurately measuring concentration for peptides with few or no chromophores(30,42) (1 Tyr residue for the peptides discussed here(9)), we used a prior of increased width for the peptide concentration, up to a limit of ±50% of the initially measured value estimated by absorbance at 280 nm. As discussed above, the posterior distributions processed through the Bayesian pipeline are limited by the most restrictive prior used, owing to the concentration ratio being well defined (S2 Table). As a result, this approach ensures that posterior distributions are limited to the range around the measured concentration of LC8, allowing us to effectively infer the uncertain peptide concentration.

**Table 1:** Ranges for thermodynamic parameters for LC8-client binding. Values delineate 95% Bayesian credibility regions from sampled posterior distributions in kcal/mol, which are akin to 95% confidence intervals. Previously published binding parameters from an identical-sites model for these isotherms are available in Supplementary Table S3.

| Peptide | $\Delta G$ | | $\Delta\Delta G$ | | $\Delta H$ | | $\Delta\Delta H$ | | $-T\Delta S$ | | $-T\Delta\Delta S$ | |
| --- | --- | --- | --- | --- | --- | --- | --- | --- | --- | --- | --- | --- |
|  | Min | Max | Min | Max | Min | Max | Min | Max | Min | Max | Min | Max |
| SPAG5 | -6.9 | -6.2 | -2.1 | -1.1 | -18 | -14 | -1.8 | 3.6 | 6.9 | 11 | -5.7 | 0.6 |
| BSN (I) | -5.6 | -4.9 | -2.1 | -0.9 | -37 | -14 | 0.1 | 45 | 8.8 | 32 | -51 | -2.6 |
| BSN (II) | -7.1 | -6.5 | -1.2 | -0.4 | -5.8 | -4.8 | -9.1 | -7.0 | -1.9 | -1.0 | 6.1 | 8.3 |
| SLC9A2 | -6.8 | -5.5 | -2.6 | -0.5 | -24 | -10 | -5.0 | 23 | 3.7 | 18.3 | -26 | 4.4 |
| VP35 | -7.4 | -6.8 | -1.7 | -0.9 | -14 | -11 | -0.7 | 1.5 | 4.0 | 6.7 | -3.0 | -0.2 |
| GLCCI | -6.1 | -5.1 | -2.6 | -1.0 | -27 | -9.7 | -0.5 | 36 | 3.7 | 22 | -39 | -0.5 |
| BIM | -9.5 | -7.1 | -2.0 | 0.8 | -12 | -10 | -0.5 | 2.9 | 1.4 | 4.4 | -0.7 | 2.2 |

Bayesian analysis of the seven systems reveals significant heterogeneity in the precision with which binding parameters can be determined (Table 1). As will be described in detail below, this is only partially reflective of apparent data quality (e.g., noise level). Instead, certain binding parameters, particularly binding enthalpies, are intrinsically more difficult to characterize. Variations in precision do not stem from inadequate sampling in the Bayesian pipeline: triplicate runs are performed to confirm sampling quality (see Methods) (example in S5 Fig.).

In particularly tractable cases, such as for SPAG5 binding in Figure 4, the analysis provides marginal distributions of similar precision to those seen with synthetic data. For binding between a peptide from the protein SPAG5 and LC8, Bayesian analysis yields a 95% credibility region of -6.9 to -6.2 kcal/mol for ΔG (Table 1), equivalent to a range for K_d1_ of 8.7 μM to 27 μM. The 95% credibility region for ΔΔG, the allosteric difference between the first and second binding event, is -2.1 to -1.1 kcal/mol, roughly equivalent to a 6 to 30-fold increase in affinity for the second binding step relative to the first. The change in binding enthalpy between first and second events, ΔΔH, is distributed around zero (Fig. 4b), with uncertainty >2 kcal/mol for all cases, meaning we are unable to discern conclusively if there is any change in enthalpy between binding steps. From ΔG and ΔH values for both binding steps, we can additionally calculate -TΔS and -TΔΔS, for the entropy of binding and the change in entropy across binding steps respectively. Although the marginal distributions for these terms are broad (Fig. 4d), the -TΔΔS mostly sits at negative values, indicating that binding enhancement has a greater probability of being entropically driven. See Table 1 for the full set of credibility regions.

**Figure 4:**
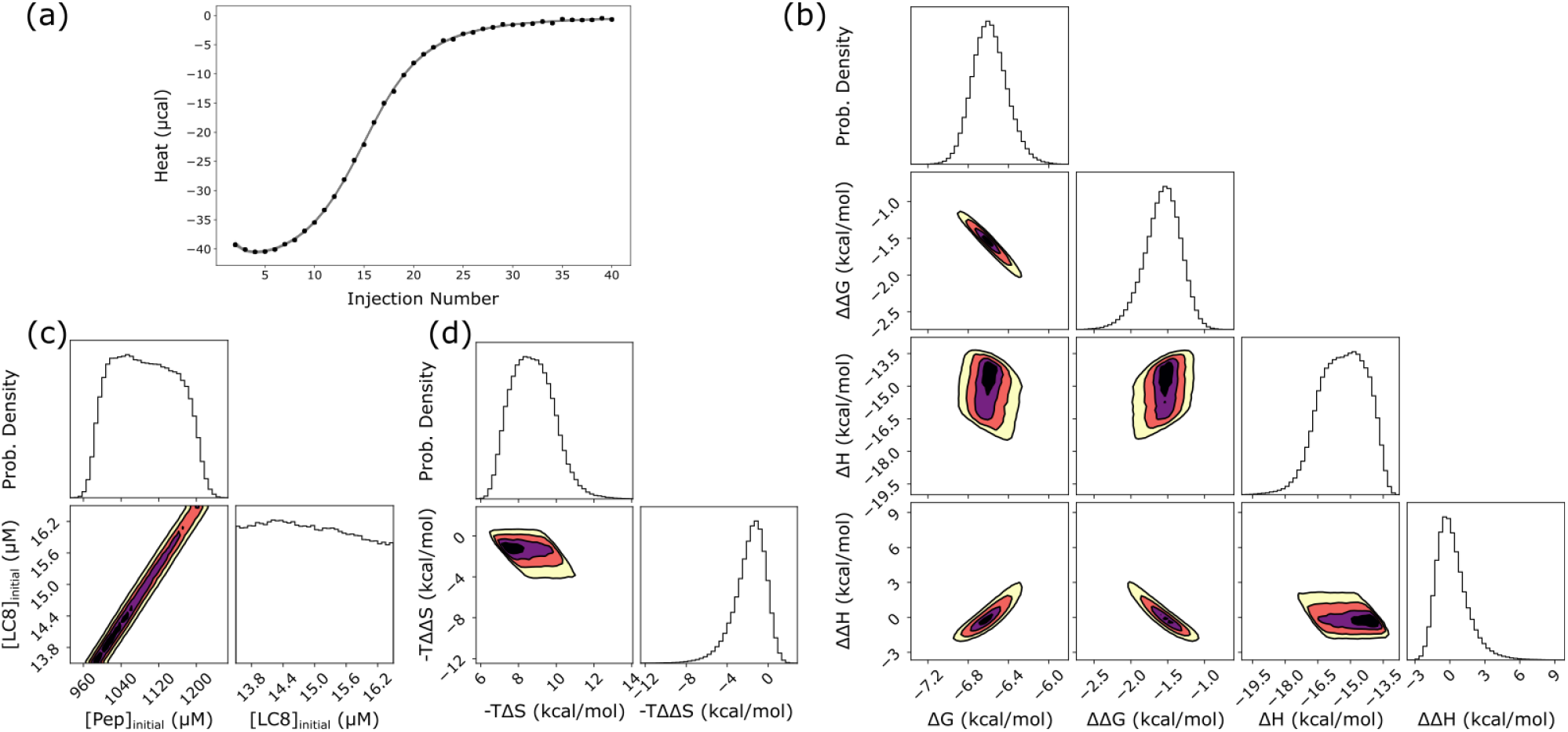
LC8 binding to a peptide from the protein SPAG5. (a) Experimental titration isotherm of SPAG5 into LC8 (points) with 50 example traces (lines) drawn from the posterior distribution of thermodynamic parameters and concentrations. (b) One and two-dimensional marginal distributions for thermodynamic parameters, with contours in the two-dimensional plots set at 95 (yellow), 75 (orange), 50(purple) and 25%(black) credibility. (c) Marginal distributions for concentrations of LC8 and peptide, showing a line of degenerate solutions, which may be compared to Fig. 2. (d) Marginal distributions for entropy (-TΔS) and change of entropy (-TΔΔS). Plots at the top of each column in panels b,c,d are one-dimensional probability density distributions.

Some general conclusions about cooperativity are apparent from the full set of data (Table 1). In all cases except one (binding to BIM), the distribution for ΔΔG is negative, indicating that all isotherms exhibit some positive cooperativity. Even for BIM, which has the widest ΔΔG distribution, the range predominantly covers negative values. All isotherms exhibit precisely determined free energies: 95% credibility regions cover a range of 2 kcal/mol or less for all cases except BIM. A common feature among some isotherms, seen clearly in the middle and right examples in Figure 5, is an apparent loss of precision in our ability to determine model enthalpies, as both show wide distributions for ΔH and ΔΔH. For these isotherms (e.g., SLC9A2, GLCCI, and BIM), the two-dimensional marginal distribution for ΔH and ΔΔH shows a clear correlative effect (S6 Fig.), and the one-dimensional distribution for the ‘total’ enthalpy (i.e. 2*ΔH* + *ΔΔH*) is narrower than the individual parameter distributions (S1 Table). In sum, the wide enthalpy distributions represent an inability to precisely determine ‘microscopic’ enthalpies for individual binding events. Nevertheless, even in these cases, the overall enthalpy can be determined with high precision (S1 Table).

**Figure 5:**
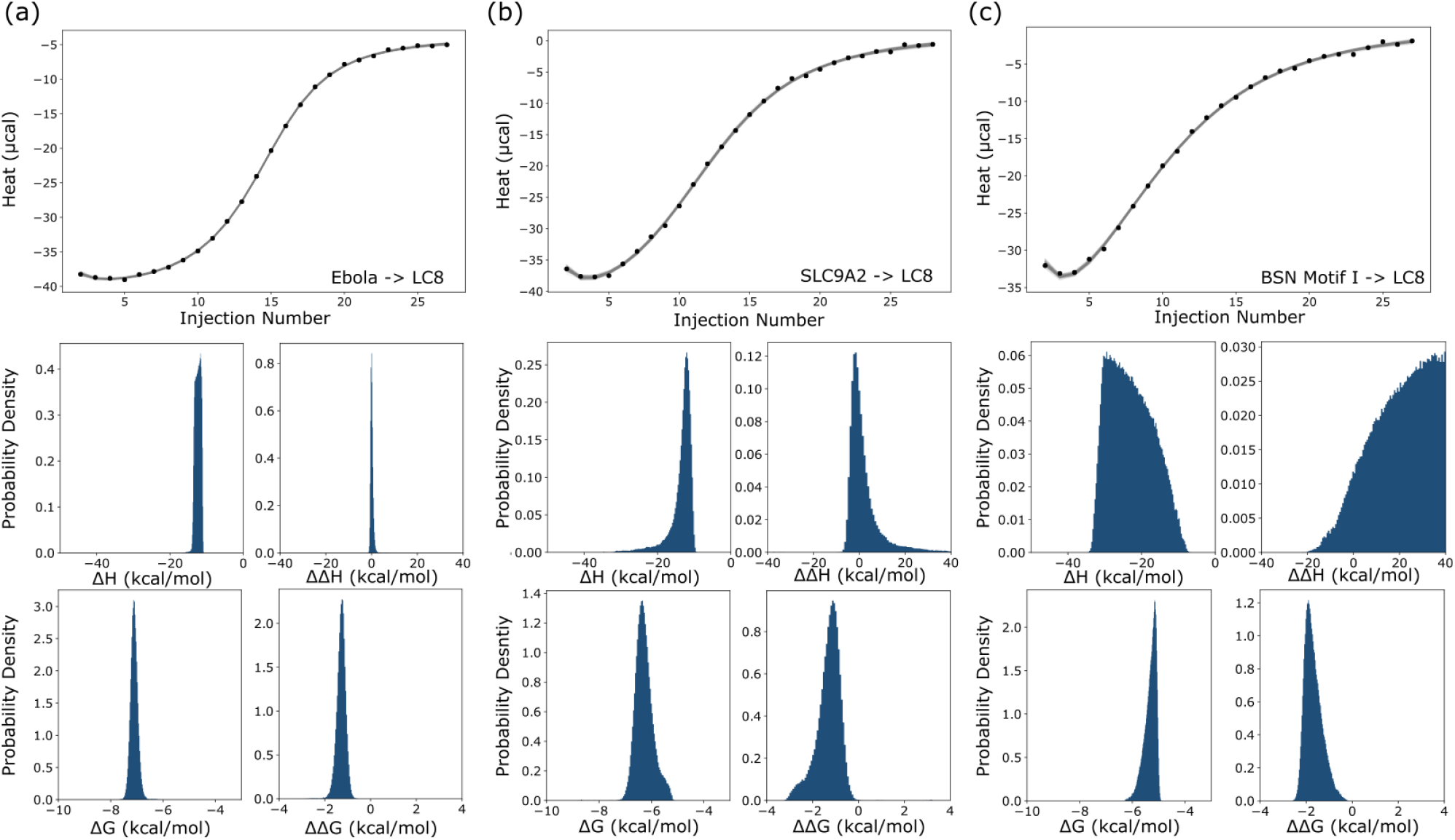
Example distributions for thermodynamic parameters from 3 LC8-peptide isotherms. Binding between LC8 and peptides from Ebola VP35 (a), SLC9A2 (b) and motif 1 from BSN (c). Isotherms are shown at the top, and distributions for thermodynamic parameters are shown below. Horizontal axes represent the full width of the uniform prior range for each parameter to allow for direct comparison between each isotherm.

### Parameter inference from multiple isotherms

The use of additional experimental information is expected to increase the precision of parameter determination, and Bayesian inference is readily adapted to employ multiple isotherms, whether at matching or different experimental conditions(38). Despite the higher dimensionality resulting from additional nuisance parameters (see Methods), we found it relatively easy to sample the parameter space for a two-site model including two-isotherms for several of our LC8-client interactions (S7 Fig.). For GLCCI, for example, the addition of a second isotherm narrowed posterior distributions, while in others (e.g. BSN motif I) it proved less impactful, largely just taking the same shape as the distribution for individual isotherms. We note that the isotherms examined were designed as technical replicates, not as optimized isotherms at different conditions for a global model. We expect results on multiple isotherms with varied experimental setups, e.g., different concentrations, to be more consistently valuable. Nevertheless, the global models demonstrate our ability to apply the pipeline to multiple isotherms simultaneously, a key step toward improved precision going forward.

### NudE-IC binding

To confirm the utility of the Bayesian pipeline for a range of proteins with two sites, we tested it on binding between the coiled-coil dimer NudE and the intermediate chain (IC) of dynein. Binding between NudE and IC can be described by the same model as binding between LC8 and clients – NudE forms a strong (K_d_ ∼ 200 nM in *C. thermophilum*(43)) dimeric coiled-coil structure which then accommodates two chains of monomeric disordered IC for a 2:2 complex stoichiometry (Fig. 6a)(39). Prior characterization of NudE-IC binding used a simple identical-sites binding model with a single K_d_, and thus provides a good system for re-analysis as well as for comparison to LC8-client binding(39,44).

**Figure 6:**
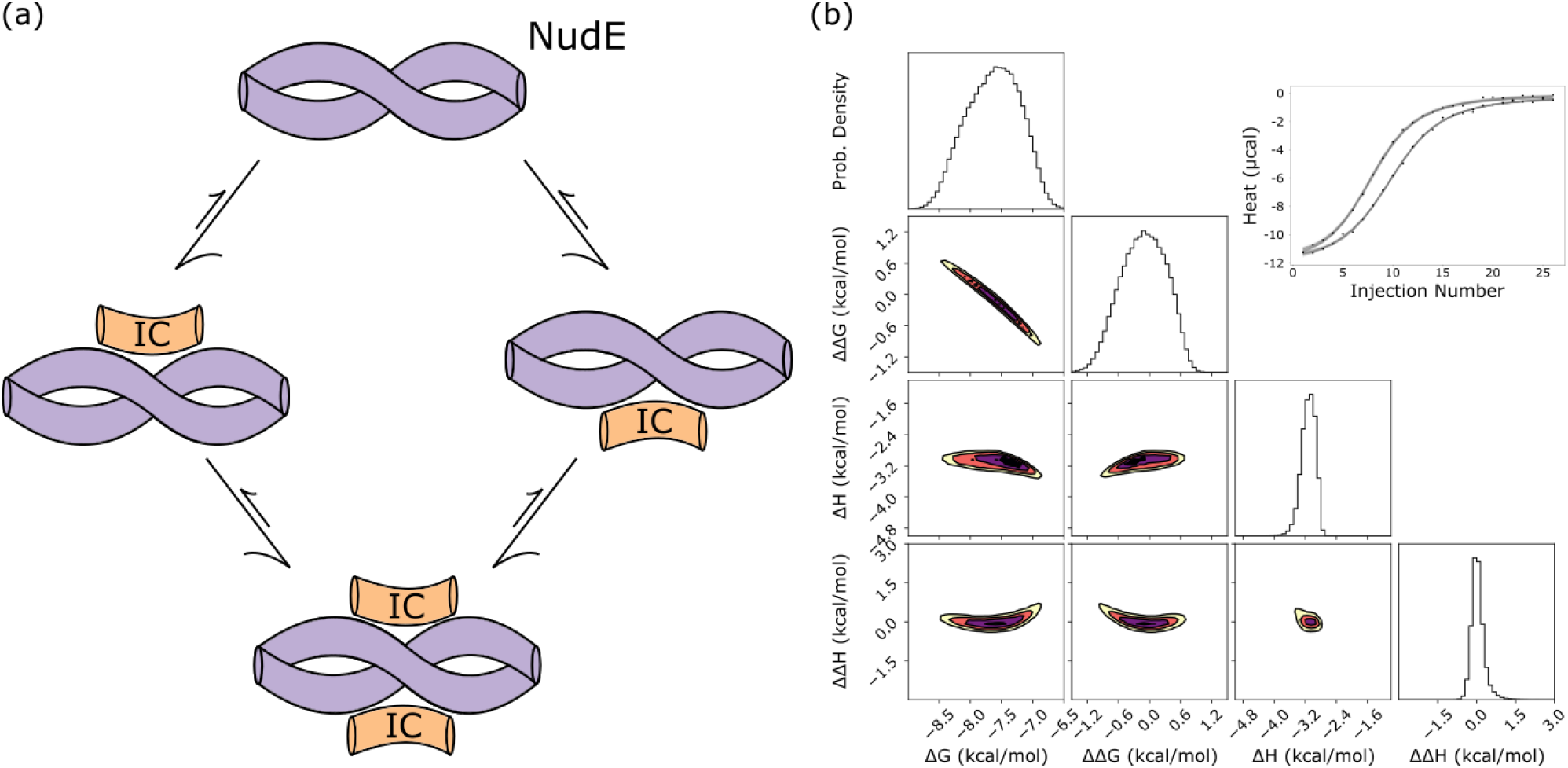
binding between the intermediate chain (IC) and NudE. (a) A model of NudE-IC binding, which forms a 2:2 complex. A cartoon diagram of NudE is shown in purple and IC in orange. (b) Sampled distributions modeled from two isotherms for binding between IC and NudE from yeast. Marginal distributions for thermodynamic parameters are shown on the left, and the top right corner contains the experimental isotherms (points) with model values (lines) drawn from the posterior.

For NudE-IC binding, a two-site model recapitulates the parameters determined in fits to identical sites modeling using Origin. For high confidence in model parameters, we applied a global model, identical to the one used in LC8-client binding, to two titrations of IC into NudE. Bayesian sampling returns narrow distributions for all thermodynamic parameters, both for individual-isotherm models (S8 Fig.), and for the global, 2-isotherm model (Fig. 6). Neither ΔΔG nor ΔΔH are significantly shifted from a distribution around zero, suggesting little, if any, cooperativity in binding. Published work applying an identical-sites model to these data provides a binding enthalpy of -3.1 kcal/mol, and an affinity of 2.3 μM (i.e. a ΔG of -7.6 kcal/mol) implying a TΔS value of 4.5 kcal/mol, meaning binding is entropically favored(45). Our two-site model predicts a ΔH distribution centered near -3 kcal/mol, and a ΔG distribution centered near -7.5 kcal/mol, aligning well with the published values. This binding interaction works well as a counterexample to LC8-client binding: distributions for allosteric terms are centered around zero and determined distributions match closely to reported values modeled from a simple model.

### Limits of precision in binding enthalpies

We exploit synthetic isotherms to systematically survey binding parameters and determine the extent to which the physical parameters themselves intrinsically lead to lower precision in parameter inference. This effort was motivated by the disparity between the three example isotherms in Fig. 5 and anecdotal observations that weaker binding, such as between LC8 and BSN I (Table 1), was correlated with increased uncertainty, i.e., broader posterior marginals, in binding parameters, especially ΔH and ΔΔH. For this purpose, we created a series of synthetic isotherms on a grid of ΔG and ΔΔG values and determined posterior distributions for each isotherm. Two-dimensional heat maps of the width of these distributions across ΔG-ΔΔG space (Fig. 7a, b) capture trends in our ability to determine model parameters.

**Figure 7:**
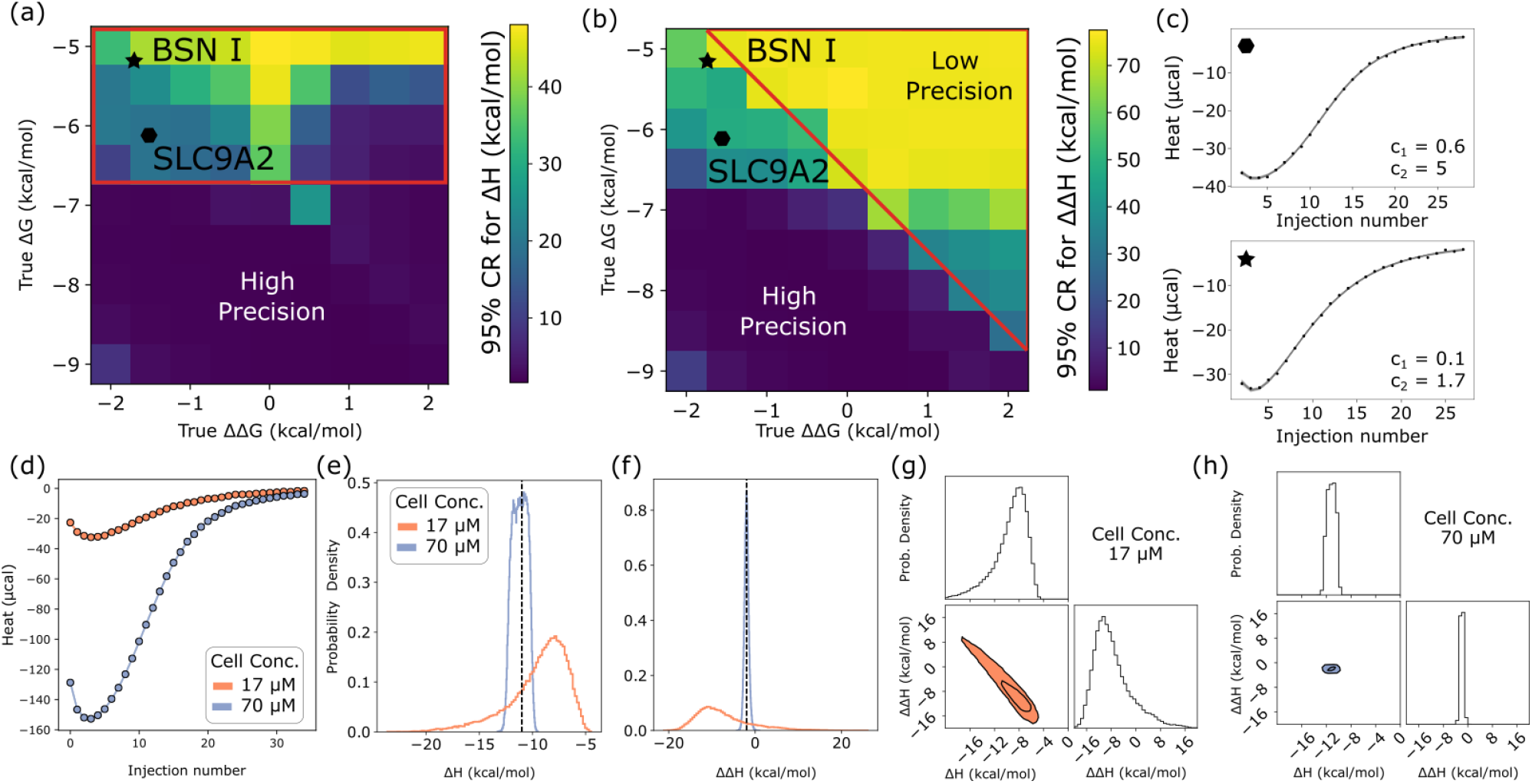
Strong dependence of posterior distributions on model parameters. (a,b) Graphical depiction of enthalpy uncertainty on a grid of ΔΔG and ΔG values, generated from the Bayesian posterior for each grid point based on synthetic data (with fixed ΔH = -10, ΔΔH = -1.5 kcal/mol). Boxes are colored by the width of the 95% credibility region for ΔH (a) and ΔΔH (b), with lighter colors corresponding to wider credibility regions (color bars). Red polygons demonstrate where each K_d_ (K_d1_ for left, K_d2_ for right) is greater than 17 μM, which is the cell concentration set for these synthetic isotherms. Black symbols indicate mean values for experimental isotherms for binding for BSN motif I (star) and SLC9A2 (octagon), for comparison. (c) Isotherms for binding between LC8 and SLC9A2 (top) and BSN I (bottom). C values for each step of binding, based on mean values taken from the posterior distribution are shown. (d) Synthetic isotherms designed to mimic BSN 1 (ΔG = -5.1, ΔΔG = -1.7, ΔH = -11, ΔΔH = -2), simulated at cell concentrations of 17 (orange) and 70 (purple) μM (syringe concentrations at 900 and 2000 μM respectively). (e,f) One-dimensional probability distributions for binding enthalpy (e) and change in enthalpy (f) for the isotherms in panel (d). (g,h) Two-dimensional marginal distributions (plotting ΔH against ΔΔH for isotherms in panel (d)). Isotherms at 17 μM in (g) and 70 μM in (h). Dimension widths are fixed for both plots for better comparison.

Generally, we lose precision in binding enthalpy when binding is weaker, although there are nuances. Interestingly, the relationship appears to differ somewhat between ΔH (Fig. 7a) and ΔΔH (Fig. 7b). For ΔH, the primary dependence appears to be on the value of ΔG, with precision decreasing when ΔΔG is 0 or negative (top right corner of 7a). However, the precision for ΔΔH depends strongly on both ΔG and ΔΔG, with the worst precision found in the top left quarter of the plot. This trend is largely consistent with a fundamental principle in calorimetric experimental design based on the experimental parameter *c*, defined as *c* = *n*[cell]/*K*_*d*_, effectively a ratio between the cell concentration and the binding affinity. While the importance of *c* is debated(46), doctrine is that 5 < *c* < 500 is required to determine binding affinities from an isotherm, as binding is either too weak or too strong for the isotherm to be information-rich outside of the 5-500 range of *c*.

Consistent with the preferred range for *c*, we begin to see losses in precision as the values of *c* for our experiments decrease. Our cell concentration for synthetic isotherms is 17 μM, meaning that *c* is ∼1 at ΔG = -6.5 kcal/mol. While *c* is primarily discussed only in the context of binding models with a single affinity, with two-site binding we can calculate two separate *c* values (*c*_*1*_ and *c*_*2*_), for the first and second step of binding respectively. Regions of the heat map with the lowest precision where *c*_*1*_ < 1 are boxed in red in the ΔH heat map (Fig 7a), and *c*_*2*_ < 1 is boxed in the ΔΔH heat map (Fig. 7b). The presence of two *c* values complicates isotherm analysis, in some cases producing isotherms that appear tight-binding by visual inspection (e.g. SLC9A2, BSN I, Fig. 7c), but have one or more values of *c* outside of the informative range, consistent with our observation that these isotherms have poorly determined binding enthalpy.

To examine if increasing *c* would improve precision, we generated synthetic isotherms to mimic BSN I, at ΔG = -5.1 and ΔΔG = -1.7 kcal/mol, and applied the model at two cell concentrations, 17 and 70 μM (Fig. 7d). Consistent with our expectations, posterior distributions for binding enthalpies narrowed dramatically (Fig. 7e,f) at the 70 μM concentration, although it is worth noting that this improvement in precision is due partly to the increased signal-to-noise ratio at higher concentrations (Supp Fig. S9). Based on these results, for isotherms like the SLC9A2 or BSN I titrations, performing additional experiments at increased analyte concentrations should resolve the poor precision in enthalpy determination. This provides an example of how microscopic binding parameters influence precision, and demonstrates that future experiments designed to investigate multisite binding will benefit from consideration of how experimental conditions relate to the energy of each step of a multistep binding interaction.

## Discussion

This work develops binding models from isothermal titration calorimetry (ITC) data for proteins with two symmetrical sites to extract thermodynamics parameters for each binding event and thus assess cooperativity. One such protein is the dimeric hub LC8, which binds over 100 client proteins at the same site. The essential role LC8 plays in regulating a variety of cell functions(5,9,47) motivates detailed mechanistic understanding of what drives recognition to its diverse partners. Using a Bayesian framework, we sought to determine precisely how much information can be extracted from a single ITC isotherm and examine how uncertainty in analyte concentration impacts model parameters, an investigation greatly aided by simulated ‘synthetic’ isotherms with known parameters. Building on prior work(32,36,38), we have advanced Bayesian analysis of binding, and applied it to rigorous biophysical characterization of LC8 dimer binding to short client peptides, as well as binding between the dimeric coiled-coil domain of NudE and the intermediate chain of dynein. We also used synthetic data to unambiguously separate effects of experimental error from system-intrinsic limitations imposed and define the limits of intrinsic (in)tractability in calorimetry.

### Bayesian inference in binding analysis, leveraging synthetic data

“How much information is contained in an ITC isotherm?” is a fundamental biophysics question that Bayesian inference is uniquely suited to answer. Building on prior work(32,36,38), we have improved the ability of the Bayesian approach to account for the intrinsic uncertainty in *both* titrant and titrand concentrations. Our approach was motivated in large part by the recognition of a mathematical “degeneracy” in ITC analysis, i.e., the existence of multiple solutions even in the absence of experimental noise, which prevents inference of a fully unique set of thermodynamic parameters. This degeneracy holds for simple 1:1 binding and apparently for arbitrary stoichiometry.

While fitting ITC data to multisite binding and other complex models is challenging, Bayesian inference yields “posterior” joint probability distributions for model parameters, providing a full description of parameter uncertainties and correlations consistent with any prior assumptions. Analogous investigation using traditional least-squares fitting commonly relies on post-analysis techniques such as the error-surface analysis implemented in SEDPHAT(21), which employs a series of maximum-likelihood fits that approximate the marginal probability distribution for a given model dimension being evaluated(37). While such analyses may be adequate in many cases, they are not integrated into many fitting pipelines, including Origin, and only rarely employed. Bayesian inference offers a direct route to uncertainties and correlations between parameters without relying on a maximum-likelihood approximation(37). The posterior distributions – the joint distribution over all binding parameters – fundamentally answer the question of the information contained in an isotherm(32,38).

Our work has benefited greatly from the use of synthetic isotherms. Built from known thermodynamic parameters, the value of synthetic isotherms as an aid in experimental design is well-established(21,32), and our Bayesian pipeline allows us to quantify the relative information content of each generated isotherm. Synthetic isotherms have allowed us to test and troubleshoot our pipeline (Fig. 3), probe the information content of isotherms under variable conditions of concentration and priors (Fig. 3, S4 Fig.), and examine how thermodynamic parameters themselves impact our ability to determine information from isotherms, resulting in the heatmap of relative tractability (Fig. 7). In the context of multi-isotherm modeling, utilizing synthetic data to design new experiments, similar to what is possible with fitting in the program SEDPHAT(21,35) will be particularly valuable.

Using synthetic data, our investigation of how concentrations impact model parameters has shown that uncertainty in concentration induces uncertainty in binding enthalpy, but has a reduced impact on free energy. This agrees well with results from Nguyen et al. (2018)(32) on 1:1 binding, indicating that the concentration-enthalpy relationship applies to all binding models. We have shown that while the individual concentrations may be indeterminable from the model alone, the ratio of concentrations can be readily determined, provided the underlying stoichiometry of binding is known (S1 Table).

From a single experimental isotherm, we sample marginal posterior distributions with widths as narrow as 1-2 kcal/mol for a two-site model with four thermodynamic parameters. Uncertainties on this scale are consistent with other Bayesian analyses(36,38). In addition to uncertainty due to experimental noise and correlative parameter relationships such as between ΔG and ΔΔG, additional uncertainty arises from our ‘skeptical’ consideration of analyte concentrations (priors of ± 10% for LC8, up to ±50% for peptides). While free energy and enthalpy parameters for individual binding steps cannot always be determined with precision, the total values accounting for both steps are less uncertain (S1 Table, S2 Fig.). When necessary, uncertainty can be reduced by careful concentration determination through multiple methods, and the use of global models derived from multiple isotherms at varied concentrations and concentration ratios

### Practical limitations of Bayesian sampling and global modeling

Bayesian statistical analysis relies on Markov chain Monte Carlo (MCMC) sampling, which requires simulating a sufficient number of steps to adequately explore the parameter space, potentially including a need to locate and sample multiple probability peaks (akin to energy basins in conformation space). For our ITC data analysis, simple MCMC sampling methods proved inadequate to sample the model space, even following sampling times of several days and over 4 million samples. While the ensemble sampler(48) used by us and others applying Bayesian models to ITC(36,38) has been robust for our purposes, adequate sampling continues to be an important limitation, especially when considering analysis of more complex models. For all work presented here, wall-clock sampling times were on the scale of minutes to hours, and hence readily feasible. More complex models could require significantly more sampling, although there is no simple scaling law that applies because of the difficult-to-predict nature of the parameter-space ‘landscape’. Global modeling of multiple isotherms may also require additional sampling: as isotherms are added to a global model, each one brings with it a new set of nuisance parameters (4 per isotherm in our work - see Methods). In our hands, global models of two isotherms could be well-sampled within half a day. While global models of technical replicates may improve signal to noise ratios, ideally, global experiments should be designed with the intent of covering several experimental conditions(36,38), and all experiments must be high quality to ensure they contribute to global fits. Efficient sampling of global models is an ongoing research direction for us.

### Cooperativity in two-site binding

Our data show that LC8 binds client proteins with positive cooperativity. Of the 7 peptides examined here, Bayesian analysis for all except one (BIM) yields a negative ΔΔG value, confirming our hypothesis that cooperativity drives LC8 binding(18) for these peptides. We additionally test binding between the intermediate chain (IC) of dynein and cargo adaptor NudE and find no evidence of cooperativity in the interaction. For the present study, we selected test LC8-binding isotherms with preference for two criteria we anticipated would leverage modeling: (1) tight-binding to LC8 and (2) an isotherm shape that breaks from a strict sigmoid. This selection process makes it difficult to say with certainty whether the cooperativity we see with LC8-client binding is unique to these clients or universal to LC8 binding. Nevertheless, we can draw some conclusions about the mechanism and function of cooperativity.

The mechanism of cooperativity in LC8-client binding appears to be entropically driven. While entropy is often the term with the widest distribution (Table 1), owing to its dependence on both the free energy and the enthalpy, there is a clear trend in our results towards positive TΔΔS values, which equates to the second binding step being more entropically favorable than the first. Relatedly, NMR dynamics measurements indicate LC8’s flexible core is rigidified on binding to clients(17,49). Since LC8-binding cooperativity necessarily requires some change in the structural ensemble of LC8, it is possible that the first binding step can be thought of as ‘paying up-front’ for the entropic cost of both binding steps–i.e., rigidifying the whole LC8 core. This mechanism would also allow for variation in cooperativity on a per-peptide basis, as the degree of rigidification in the core seen by NMR is dependent on client sequence(17). Future molecular dynamics simulations can examine the differences in rigidity of the LC8 core in different bound states and across binding to different peptides.

While LC8-client complexes are varied, the putative functional unit of most LC8-client interactions is a 2:2 bound structure, where LC8 promotes dimerization in client proteins(50–52). One possible underlying function of cooperative binding is therefore that it acts as a driver of the formation of the 2:2 bound state and suppresses the nonfunctional 1:2 intermediate. LC8 contrasts interestingly with NudE-IC binding in this respect. While our isotherm examines the dimerization of IC by NudE, it is unlikely that NudE ever dimerizes IC in biological conditions. In fact, recent work has demonstrated that pre-dimerization is essential for the NudE-IC interaction when measured in context of the full length protein(43). As such, there is no analogous functional role for cooperativity to play in IC binding. Prior work has proposed that positive cooperativity could drive the formation of homologous complexes(18), with the same client bound to each LC8 motif. This hypothesis suggests that binding at one site essentially promotes binding of the same client at the second site, through small adjustments along the LC8 dimer interface. Future competition assays – titrations of a client into LC8 samples pre-bound with a different client – may help to answer this question.

New questions arise when considering LC8 clients with multiple binding motifs. Proteins such as the nucleoporin Nup159 or the transcription factor ASCIZ contain several LC8 motifs in succession and form large ladder-like complexes with LC8. Complexes between LC8 and multivalent clients are often highly heterogeneous in both stoichiometry and conformation(7,53,54), complicating their analysis. Particularly in the case of ASCIZ, the fully bound state is highly disfavored by negative cooperativity(7,54), thought to be mediated through the linker sequences between LC8 motif, which ensures that ASCIZ is sensitive to LC8 even at high LC8 concentrations. Investigating the thermodynamics of such complexes necessitates methods able to dissect the complicated network of binding interactions dependent both on the individual motifs as well as on the lengths and structures of linkers between motifs. This work represents a first step towards such investigations and lays a framework upon which analyses of multivalent LC8 interactions can be built.

### Concluding remarks and future steps

Bayesian inference has allowed us to characterize multi-site binding cooperativity with high confidence for two different protein-protein interactions of 2:2 stoichiometry, despite uncertainties in analyte concentrations and inherent limitations of ITC. Our analysis was enabled by improvements to prior work(32,36,38) in treating concentration uncertainties, and further demonstrates the value of Bayesian inference to ITC analysis. We used synthetic data to systematically characterize the uncertainty landscape for 2:2 binding based on intrinsic binding properties, an approach that can readily be extended to other models.

We examined two dimeric systems, the hub protein LC8 and the coiled coil domain of NudE. For LC8, every client peptide studied showed evidence of cooperative binding, confirming hypotheses from a decade ago(18). In contrast, the dynein NudE/IC complex showed minimal evidence of cooperativity, consistent with the fact that in biological settings, NudE binds to IC in a dimeric state, suggesting allostery would serve no purpose in the interaction. The ability to reliably characterize these interactions also serves as an important step toward quantitative characterization of multivalent LC8-multivalent client complexes, which, due to their complexity, evade straightforward investigation.

While our focus here has been on two-step symmetric-site binding systems, Bayesian methods can be applied to other complex models investigated by ITC. Measurement of complex multivalent systems, enthalpy-entropy compensation, and ternary complexes or competition binding are all likely to benefit from analysis under a Bayesian framework. Although there is a limit on how much information can be gained from individual isotherms, investigation utilizing synthetic data can guide design, to help determine experimental conditions that maximize gain from additional ITC experiments within a given system.

## Author Contributions

A.B.E. – Performed research, assisted in method design and development, curated and analyzed data, wrote the manuscript

A.G. – Designed and developed modeling method, assisted in writing the manuscript

E.J.B. – Designed research, Edited the manuscript

D.M.Z. -- Designed research, supervised method design and development, Edited the manuscript

## Methods

### 1:1 Binding

For 1:1 binding, we used the quadratic model as described in equations 1 and 2 in the results above. The heat of each injection *i*, (*dQ*_*i*_) was calculated using the following equation:

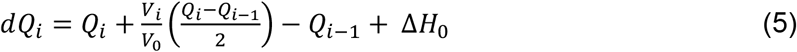

Where *V*_*i*_ is the volume of injection *i*. This binding model is identical to the model used in Origin’s identical sites model when *n* = 1. Δ*H*_0_ is a correction term to account for heat of dilution, buffer mismatch, and other effects that may apply a flat shift to binding heat.

### Two-site binding

Two-site binding is modeled in a standard fashion, such as in the binding polynomial model (55) as:

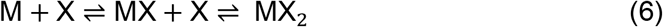

Under this scheme, each binding affinity is as follows:

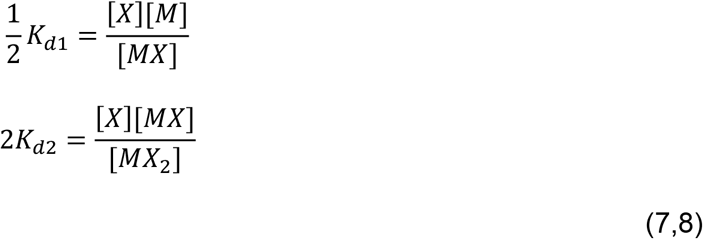

where K_d1_ and K_d2_ are the affinities for the first and second microscopic, site-resolved binding steps. Factors of ½ and 2 represent adjustments between the individual site-resolved affinities for either symmetric indistinguishable binding step and the ‘macroscopic’ binding constants (i.e. *K*_*macro*1_ = ½ *K*_*d*1_). Here and throughout, we refer to binding affinities and energies by their microscopic value. The total concentrations of X and M can be written as:

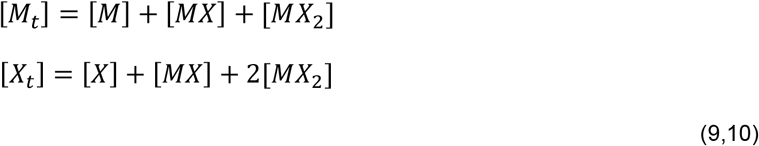

Through rearrangement and substitution of equations 7 and 8, the total concentration equations can be rewritten only in terms of [M] and [X], the concentrations of free macromolecule and ligand:

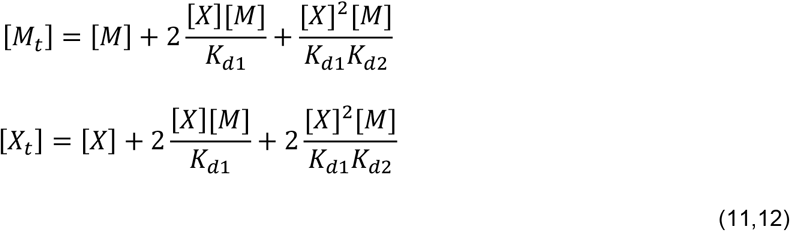

This system of equations is solved numerically for each given injection point to determine the unbound concentrations [M] and [X]. In our implementation, numerical solutions are calculated using the ‘oprimize.root’ function in the scipy python library, using a Levenberg-Marquardt least-squares optimization. With both free concentrations determined, the system heat can be calculated:

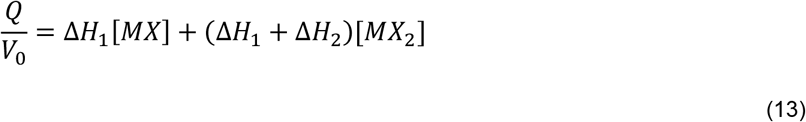

where ΔH_1_ and ΔH_2_ are the enthalpies of binding step one and two respectively. The concentrations of each bound state can be calculated from [X] and [M] and equations 7 and 8. As in the 1:1 binding model, equation 5 is used to calculate the observed heat of injection, *dQ*_*i*_, for each injection.

### Degeneracy in two-step binding

When protein concentrations are included as model parameters, degenerate solutions are introduced. As outlined in the manuscript, the degeneracy is exposed from the following transformation:

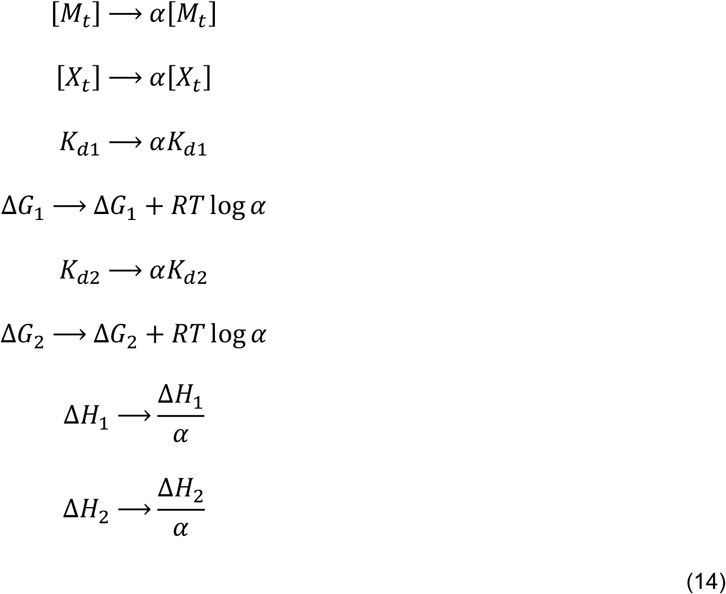

Here, α can be any positive number. Following this transformation, the equations used to calculate [X] and [M] (eq. 7 and 8) are transformed:

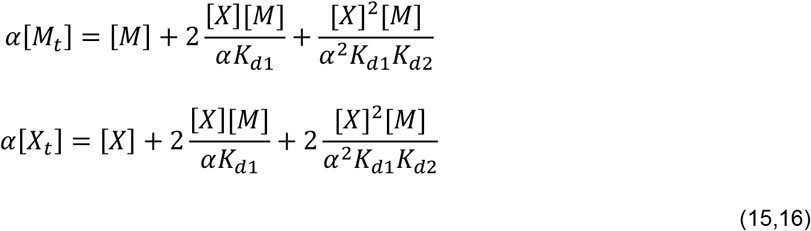

In these transformed concentration-sum equations, the new solutions for both [X] and [M] are exactly the previous solutions multiplied by *α*, as can be verified by substitution. Finally, applying the transformed values into the equation for Q yields

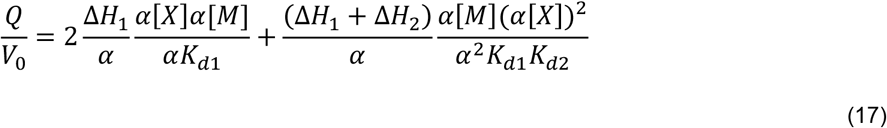

As in the 1:1 binding model, cancellation of α shows there is no change in the value of Q for any α value. This demonstrates the degeneracy for 2:2 binding, which we can expect to generalize to higher stoichiometries.

### Bayesian inference

Bayesian inference is a method to calculate a “posterior” *distribution* of model parameter values based on prior assumptions (encoded as prior distributions for parameters presumed to hold in the absence of data) and the data. In general, as more data is analyzed, the influence of the prior will decrease(56,57). The posterior distribution of parameters provides rich information such as the parameter means and confidence intervals (technically “credibility regions”), in addition to correlation information regarding whether and how parameters vary together.

Bayesian inference is based on Bayes’ rule(56,58) which enables us to infer a distribution of parameters θ (e.g., binding free energy and enthalpy, etc.) consistent with a given set of data D (e.g., ITC isotherms):

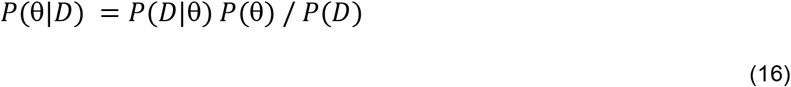

where *P*(θ|*D*) is the (posterior) probability distribution of the model parameters, θ, given the data, D; *P*(*D*|θ) (the likelihood) is the probability distribution of the data given the model parameters and is given below; *P*(θ) (the prior) is the probability of the model parameters, specified below; and *P*(*D*) (the evidence) is the probability of the data. For a given set of data, the unknown denominator *P*(*D*) is constant, independent of parameters, so it does not affect the inference of posteriors. Typically, it is not possible to analytically solve Bayes’ rule, so numerical methods such as Markov chain Monte Carlo are used to determine the target (posterior) distribution(59–61). Details of our implementation are given below.

### Bayesian model

Following prior work(30,38), we assume the data has Gaussian noise with a mean of zero and an unknown standard deviation. The ITC model parameters θ include concentration terms (X_initial_,M_initial_) and thermodynamic terms (ΔG, ΔΔG, ΔH, ΔΔH), as well as the nuisance parameters (ΔH_0_ and σ) for heat of dilution and Gaussian noise. We use uniform prior distributions for the model parameters specified below and the unknown noise standard deviation unless otherwise stated. For global models (e.g. S7 Fig., Fig. 6), while it may be possible to assume a global noise or concentration model, we instead elected to apply global models with an additional set of concentration and nuisance parameters for each additional isotherm (bringing the total parameter count up to 12 for two-isotherm models). Uniform prior ranges for thermodynamic parameters were identical for all models, listed in supplemental table S4. For nuisance parameters ΔH_0_ and σ, uniform priors of -10 to 10 μcal and 0.001 to 1 μcal respectively were used in all models.

The likelihood for a set of data *D* = {*x*_1_, *x*_2_, … }, denoted (*p*(*D*|*θ*)), is the product of the probabilities at all data points *x*_*i*_ based on a normal distribution of standard deviation σ centered around *μ*_*i*_(*θ*), the calculated value of point *i* for the binding model and parameters θ. It therefore takes the following form:

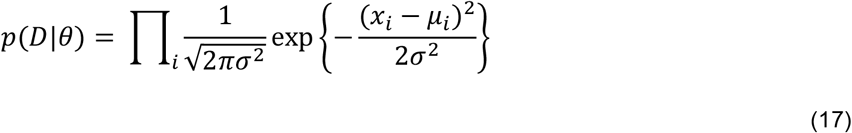

and we note that σ is assumed unknown and sampled as part of the Bayesian inference process. When the priors are uniform, as we most often assume, the posterior is simply proportional to the likelihood given here.

### Sampling

We use an affine-invariant Markov chain Monte Carlo sampling method(62) to perform Bayesian inference, as also used by Duvvuri et al. (2018)(38) and Cardoso et al. (2020)(36). The affine-invariant sampler is an ensemble-based method in which multiple walkers move through the sample space in a correlated fashion. We empirically found this method to sample significantly better than the standard Metropolis-Hastings (59,60) sampler for our model. In our hands, the Metropolis-Hastings method was unable to converge on the target distribution after 4,000,000 sampling steps, whereas the affine-invariant sampler was able to converge after 100,000 sampling steps.

### Implementation

We used the EMCEE package(48) in Python to perform the affine sampling, using a 20%:80% mix of the “differential evolution” and “stretch” move sets with 25-50 walkers. For each experiment, 3 replicas are run for 50,000-200,000 sampling steps/replica until convergence. Each replica converged, as determined by the autocorrelation time, where sampled steps must be greater than 50x the autocorrelation. Convergence was additionally assessed through examination of posterior distributions from model replicas, which were nearly identical in all cases (S5 Fig.). This implementation runs at ∼9 samples for each walker per second on 4 cores of a node on the Oregon State College of Science computing cluster.

The code, data, and an example notebook are available at: https://github.com/ZuckermanLab/Bayesian_ITC

### Synthetic isotherms

Synthetic isotherms for 1:1 and two-site binding were generated following equation 2 for 1:1 binding and equations 11, 12, and 10 for two-site binding. Parameters were chosen to mimic typical experimental conditions employed in our group. For 1:1 binding (S4 Fig.), we used ΔG and ΔH values of -8 and -12 kcal/mol respectively, and concentrations of 34 μM in the cell and 500 μM in the syringe. For two-site binding, varied thermodynamic parameters were used (e.g. Fig. 3, Fig. 7), but concentrations were fixed at 17 μM in the cell and 500 μM in the syringe. Synthetic isotherms used a cell volume of 1.42 mL and a temperature of 25 C. For synthetic isotherms, we simulated one injection of 2μL followed by 34 injections of either 6 μL (Isotherms in Fig. 3, S1,S3,S4 Figs.) or 10 μL (BSN mimic isotherms, Fig. 7d-h, S9 Fig.). All isotherms were calculated with a ΔH_0_ of 0 μcal, and added synthetic noise from a Gaussian distribution with a mean of 0 and standard deviation 0.2 μcal (except the high noise BSN I isotherm, S9 Fig.). To accurately replicate experimental conditions, we eliminated the first injection when applying models to this data.

## Acknowledgements

This work was supported by the U.S. National Institutes of Health grant R01-GM141733 and by U.S. National Science Foundation Grant MCB 2119837. We additionally acknowledge the support in the form of computational resources of the Oregon State University NMR Facility funded in part by the National Institutes of Health, HEI Grant 1S10OD018518, and by the M. J. Murdock Charitable Trust grant #2014162. We appreciate helpful discussions with John Chodera, David Minh, and Trung Hai Nguyen.

