## Supplementary Info for "Quantifying cooperative multisite binding in the hub protein LC8 through Bayesian inference"

### Contents

**Figures:**

**S1 Figure:** example MCMC traces and marginal distribution for all model parameters.

**S2 Figure:** Distributions of thermodynamic parameters plotted with total free energies and enthalpies.

**S3 Figure:** Marginal distributions comparing models with uniform and normal-distribution priors.

**S4 Figure :** Effect of concentration priors on marginal posterior distributions for thermodynamic parameters in a 1:1 binding model.

**S5 Figure :** Example marginal distributions of replicate models for the LC8-SPAG5 interaction.

**S6 Figure :** two dimensional marginal distributions of enthalpy for selected isotherms.

**S7 Figure :** Marginal distributions for thermodynamic parameters for individual and global models for three LC8-peptide interactions.

**S8 Figure :** Marginal distributions for thermodynamic parameters for IC-NudE binding isotherms.

**S9 Figure :** Marginal distributions for thermodynamic parameters for BSN-like synthetic isotherms

**S10 Figure :** Direction of injection impacts parameter determinability.

**Tables:**

**S1 Table:** Credibility regions for ‘sum’ thermodynamic parameters and ratios of concentrations.

**S2 Table:** Ranges for thermodynamic parameters for LC8-client binding when modeled with ±20% LC8 concentration.

**S3 Table:** Binding parameters determined from identical sites model fits, as published in Jespersen et. al. (2020).

**S4 Table:** Model priors and sampling lengths for all isotherms.

**Documents:**

**S1 Document:** Best practices for the application of Bayesian statistical models to isothermal titration calorimetry

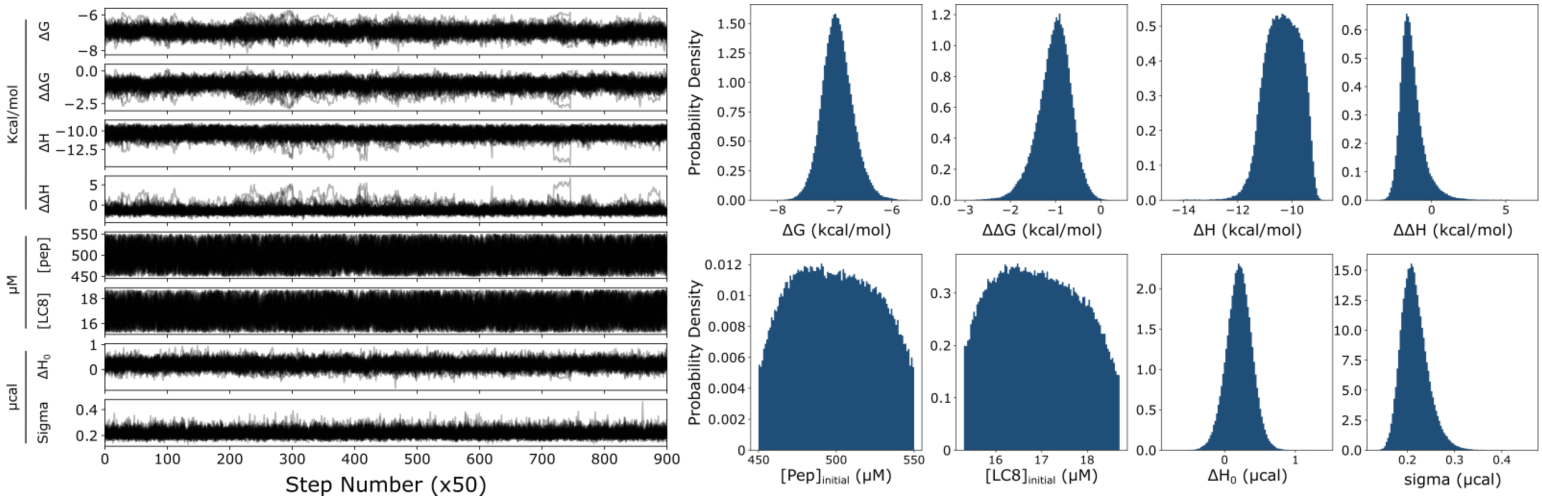


**S1 Figure: example MCMC traces and marginal distribution for all model parameters.** MCMC traces and distributions are drawn from model on synthetic isotherm from fig. 3, generated from parameters $\Delta G = -7$, $\Delta\Delta G = -1$, $\Delta H = -10$, $\Delta\Delta H = -1.5$, $[\text{peptide}]_{initial} = 500$, $[\text{LC8}]_{initial} = 17$, $\Delta H_0 = 0$ and sigma = 0.2. Each trace includes multiple walkers, each of which is drawn as its own chain (see methods for details). Traces are thinned to one step for every fifty.

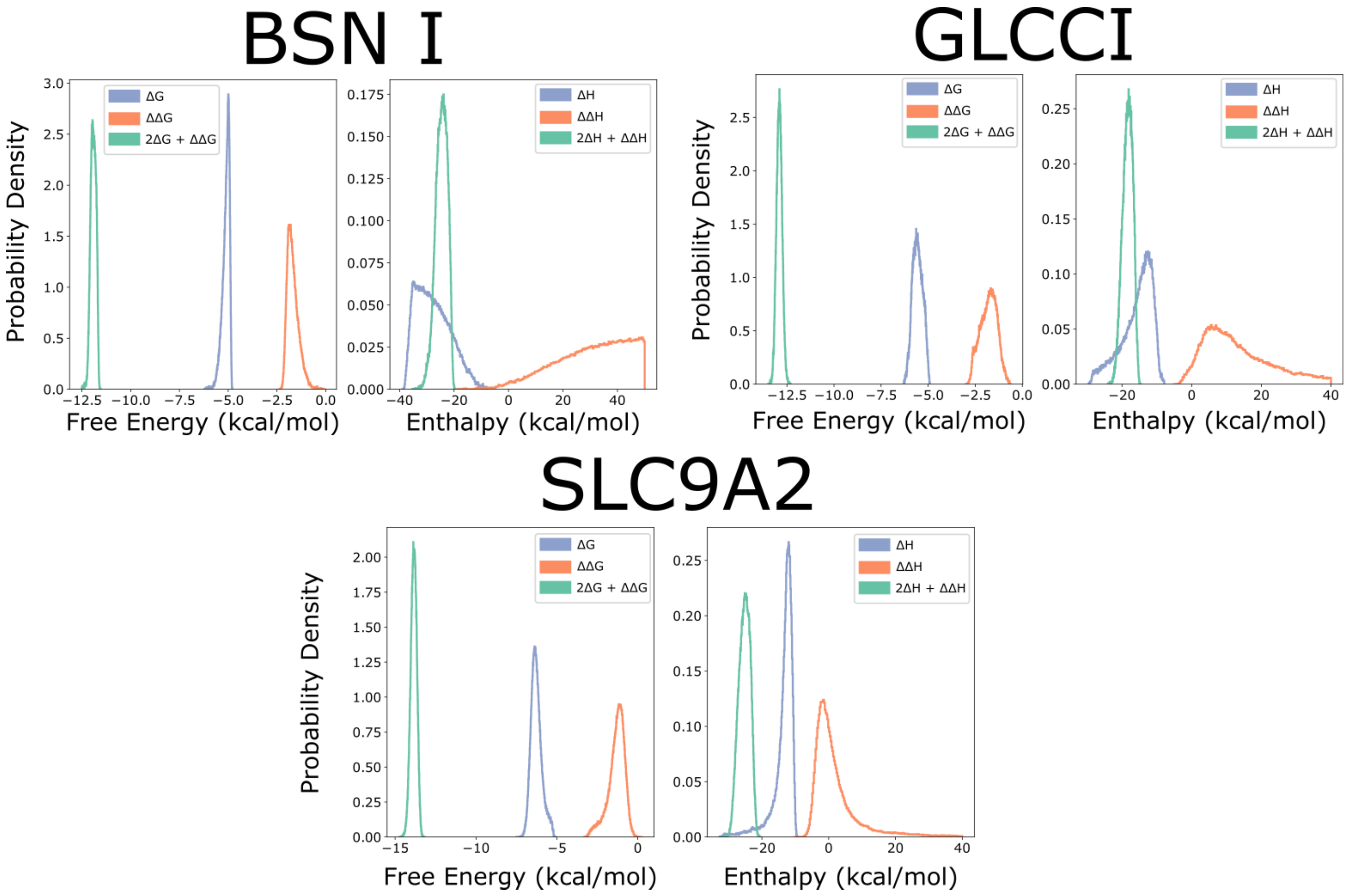


**S2 Figure: Distributions of thermodynamic parameters plotted with total free energies and enthalpies.** Each plot shows a set of either $\Delta G$ and $\Delta\Delta G$ or $\Delta H$ and $\Delta\Delta H$, along with the 'total' value for that parameter, i.e. $2\Delta G+\Delta\Delta G$ or $2\Delta H+\Delta\Delta H$. The distributions for this sum value are often narrower than the individual parameters, as the total enthalpy and free energy of binding can be determined with higher precision from a given isotherm than the individual values. $\Delta G$,$\Delta H$ are the energy and enthalpy of binding step 1, while $\Delta G+\Delta\Delta G$,$\Delta H+\Delta\Delta H$ are the energy and enthalpy of binding step 2, making the total values reported here the energy and enthalpy of both binding steps combined.

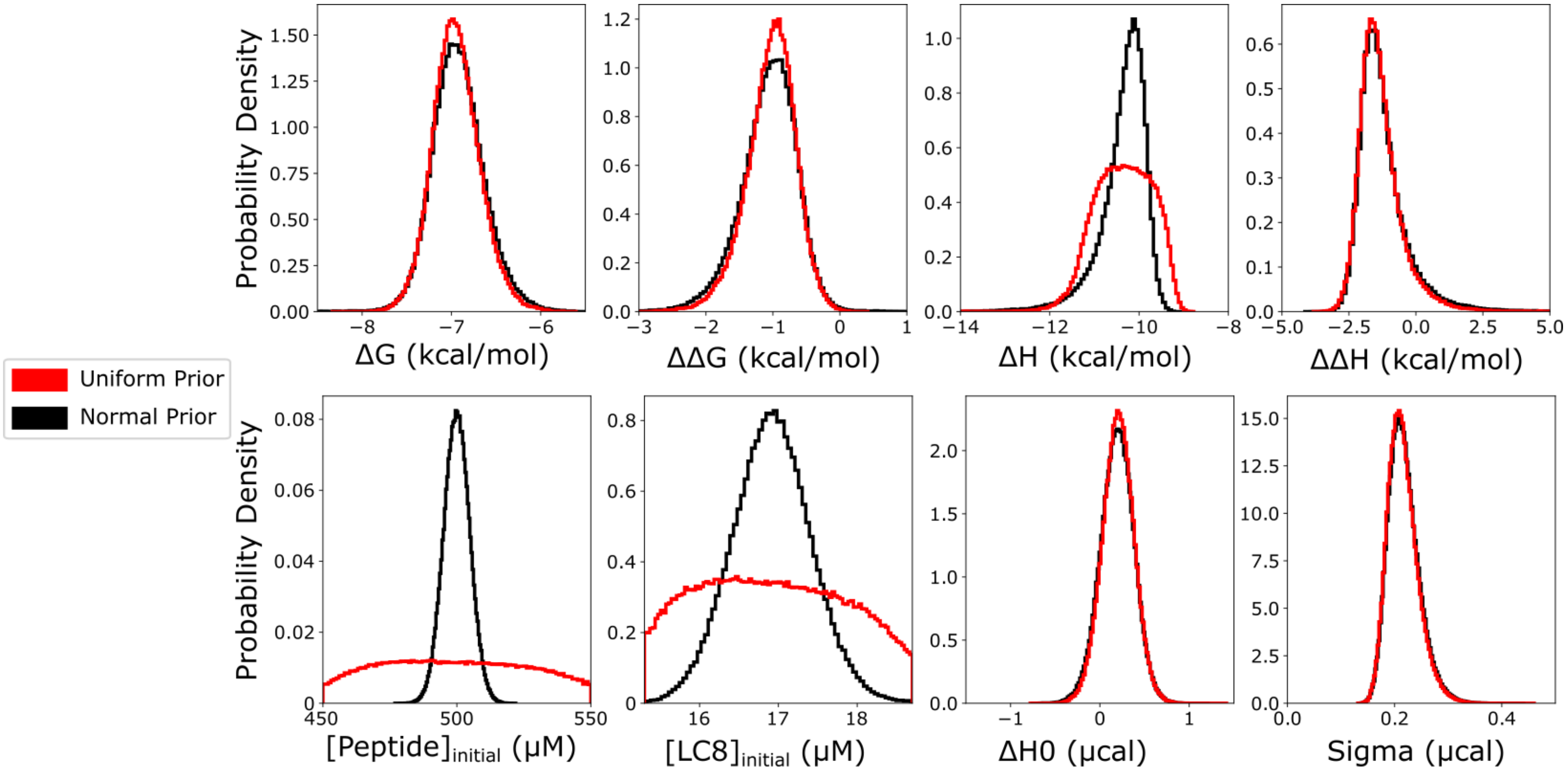


**S3 Figure: Marginal distributions comparing models with uniform and normal-distribution priors.** Distributions are taken from models on an identical synthetic isotherm generated from parameters ΔG = -7, ΔΔG = -1, ΔH = -10, ΔΔH = -1.5, $[\text{peptide}]_{initial}$ = 500, $[\text{LC8}]_{initial}$ = 17, $\Delta H_0$ = 0 and sigma = 0.2. All priors are identical except for [peptide]initial, where the uniform prior model (red) was run with a ±10% of stated value uniform prior, and the normal prior model (black) was run with a normal distribution prior with standard deviation = 1% of stated value.

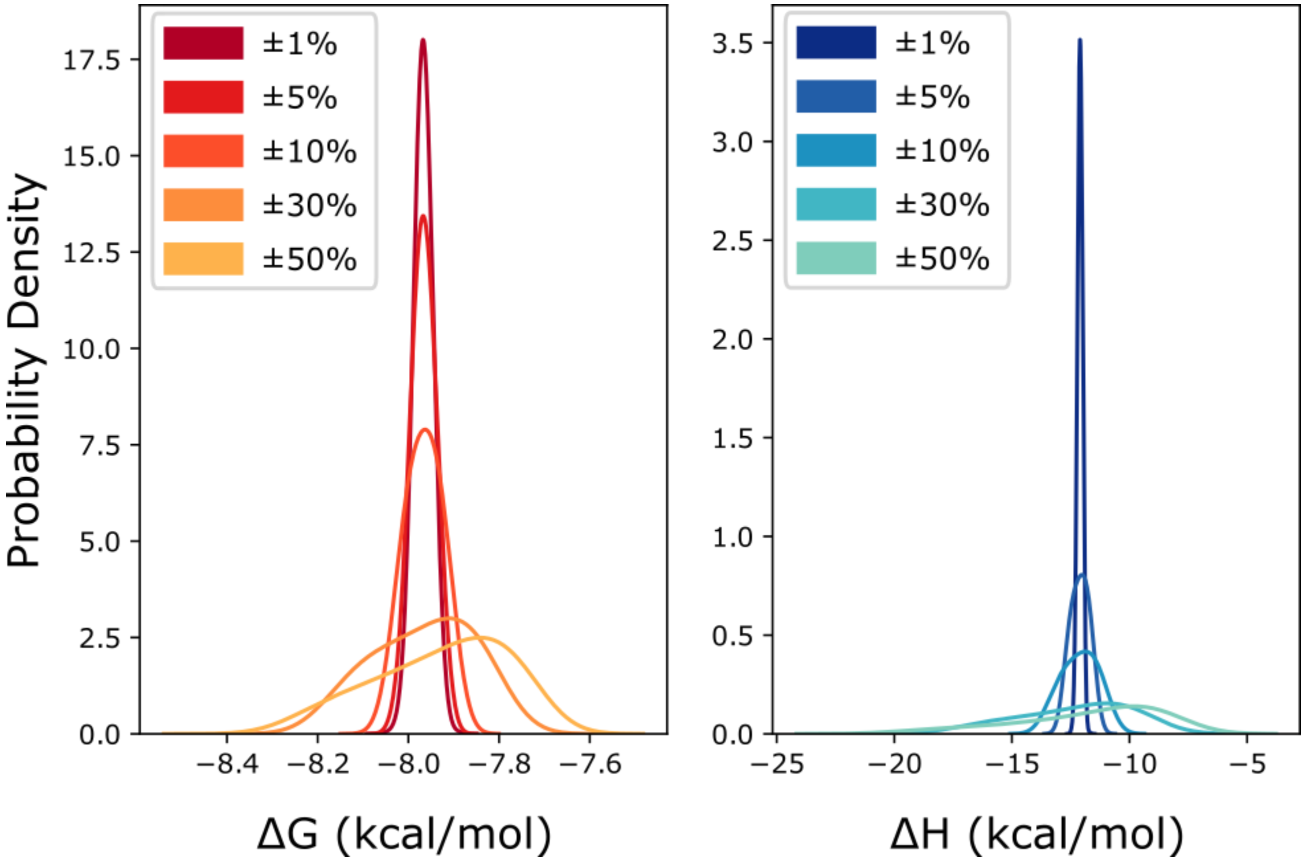


**S4 Figure: Effect of concentration priors on marginal posterior distributions for thermodynamic parameters in a 1:1 binding model.** Distributions are taken from models on an identical synthetic isotherm generated from parameters $\Delta$G = -8, $\Delta$H = -12, $[X]_{initial}$ = 500, $[M]_{initial}$ = 34, $\Delta H_0$ = 0 and sigma = 0.2. Model priors ar0e uniform distributions of varied width in each plot for $[X]_{initial}$ and $[M]_{initial,}$ varied from ±1% to ±50%.

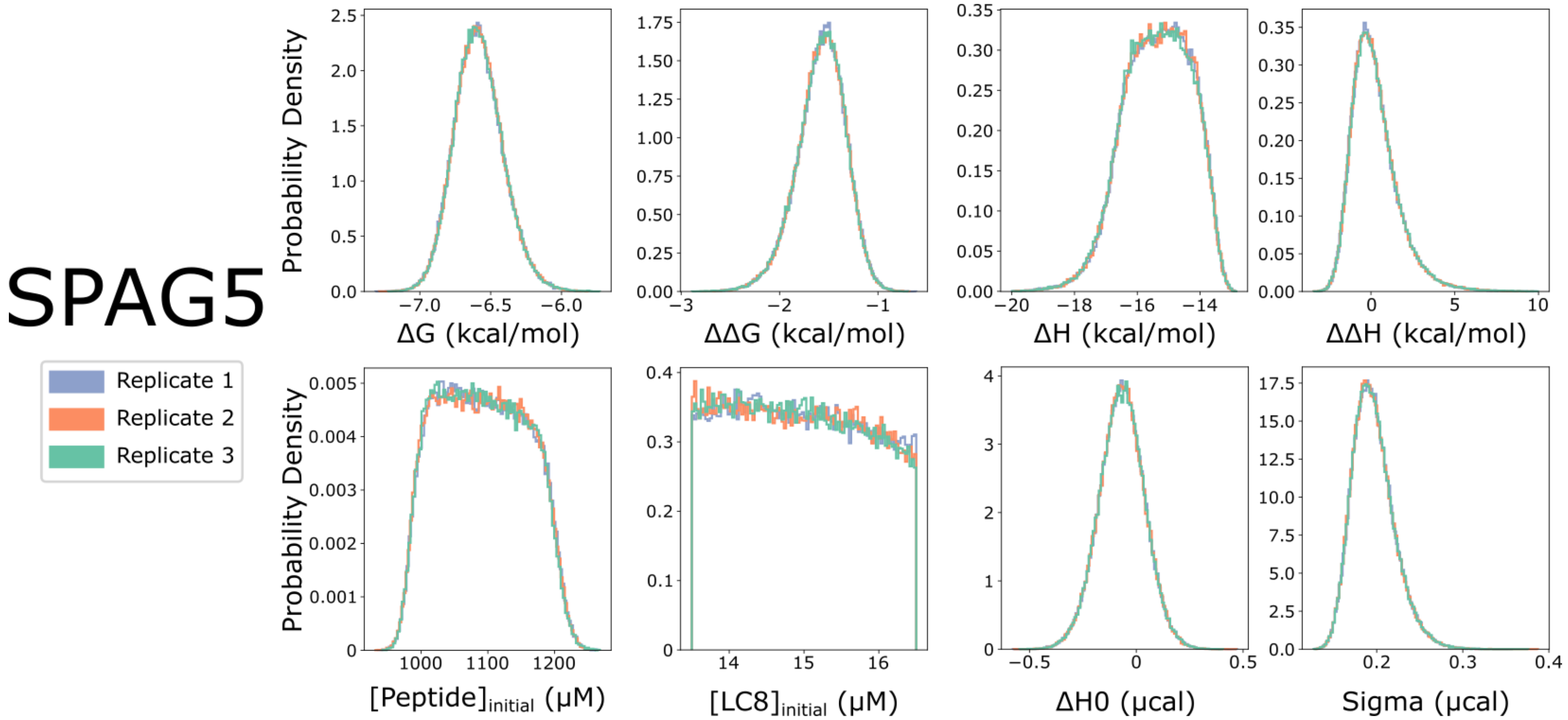


**S5 Figure: Example marginal distributions of replicate models for the LC8-SPAG5 interaction.** Each model replicate is run on an identical isotherm with a different random seed dictating random starts for MCMC chains and trial move selections. Each model returns near-identical marginal distributions.

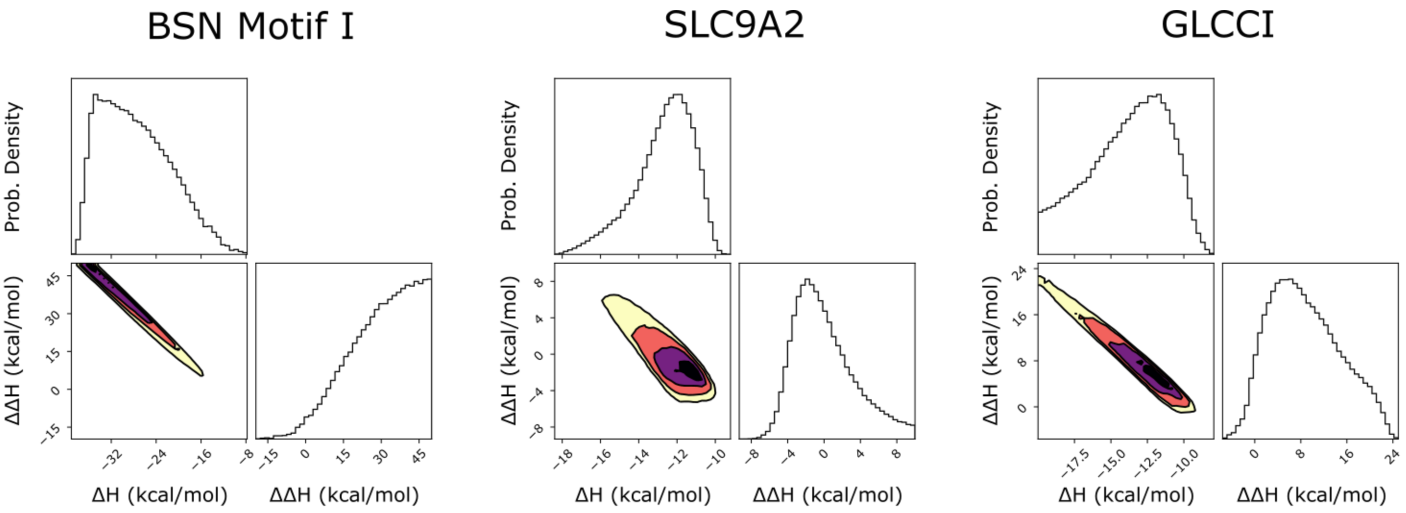


**S6 Figure: two dimensional marginal distributions of enthalpy for selected isotherms.** Marginal distributions for BSN I, SLC9A2, and GLCCI are shown, each of which has wide 1D distributions for both $\Delta$H and $\Delta\Delta$H. Enthalpy parameters are closely correlated, resulting in a diagonal two-dimensional distribution within the enthalpy space.

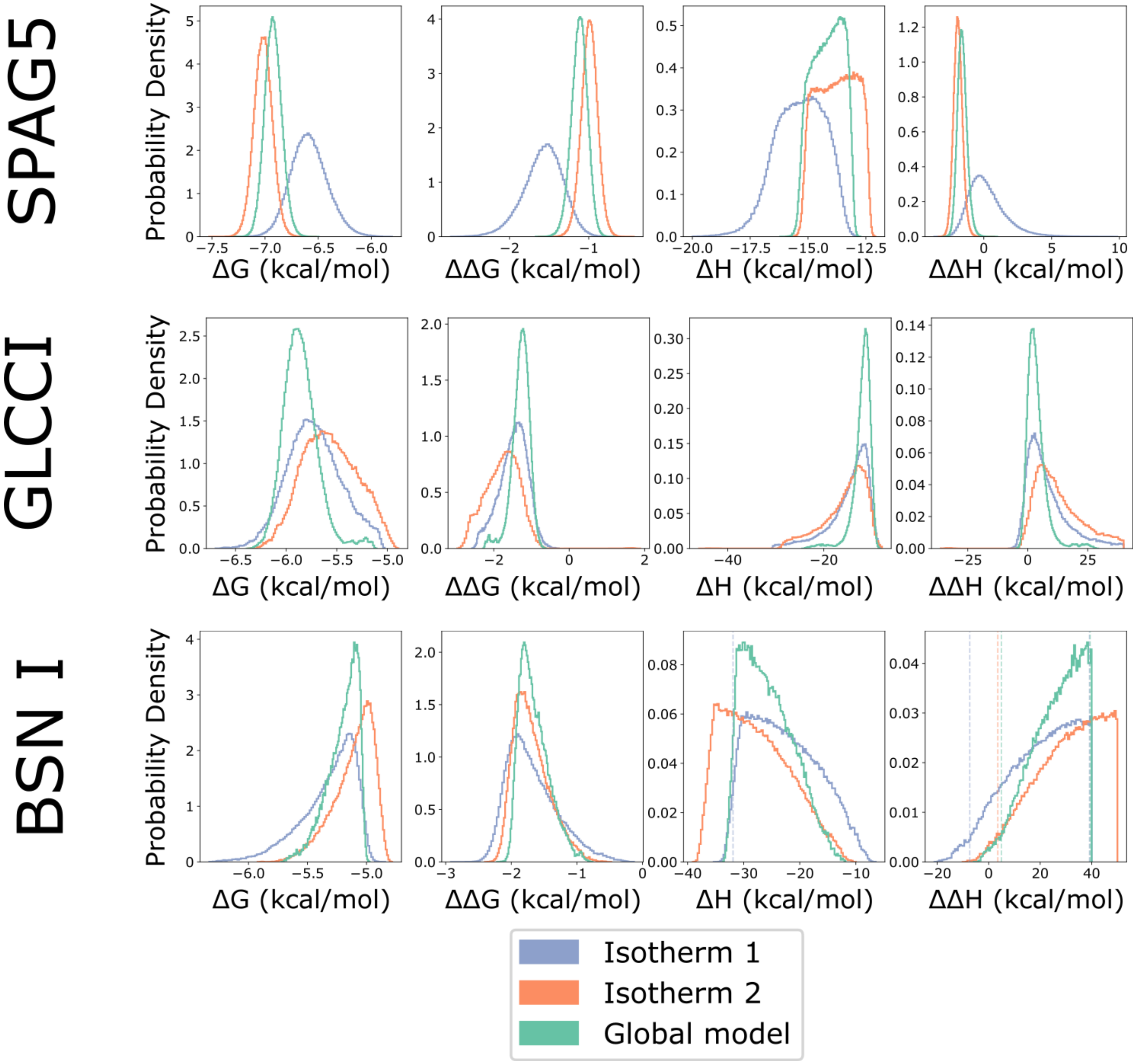


**S7 Figure: Marginal distributions for thermodynamic parameters for individual and global models for three LC8-peptide interactions.** Distributions for each individual isotherm and distributions for the global model are shown in purple, orange and green respectively. While the global model improves precision in determined parameters in some cases (e.g. GLCCI), in others it appears to follow the shape of the distributions for individual isotherms (e.g. BSN I).

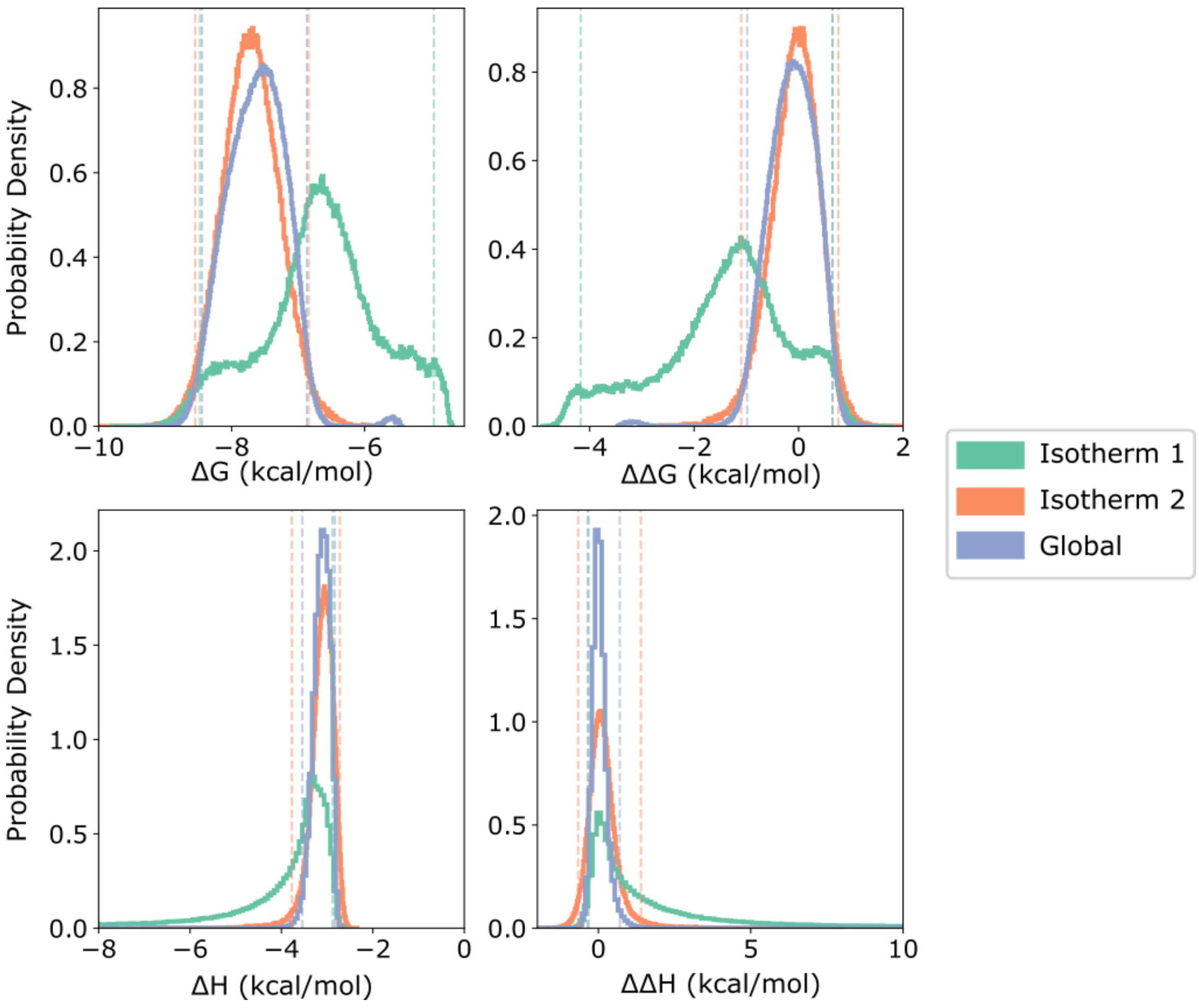


**S8 Figure: Marginal distributions for thermodynamic parameters for IC-NudE binding isotherms.** Distributions for each individual isotherm and distributions for the global model are shown in green, orange, and purple respectively.

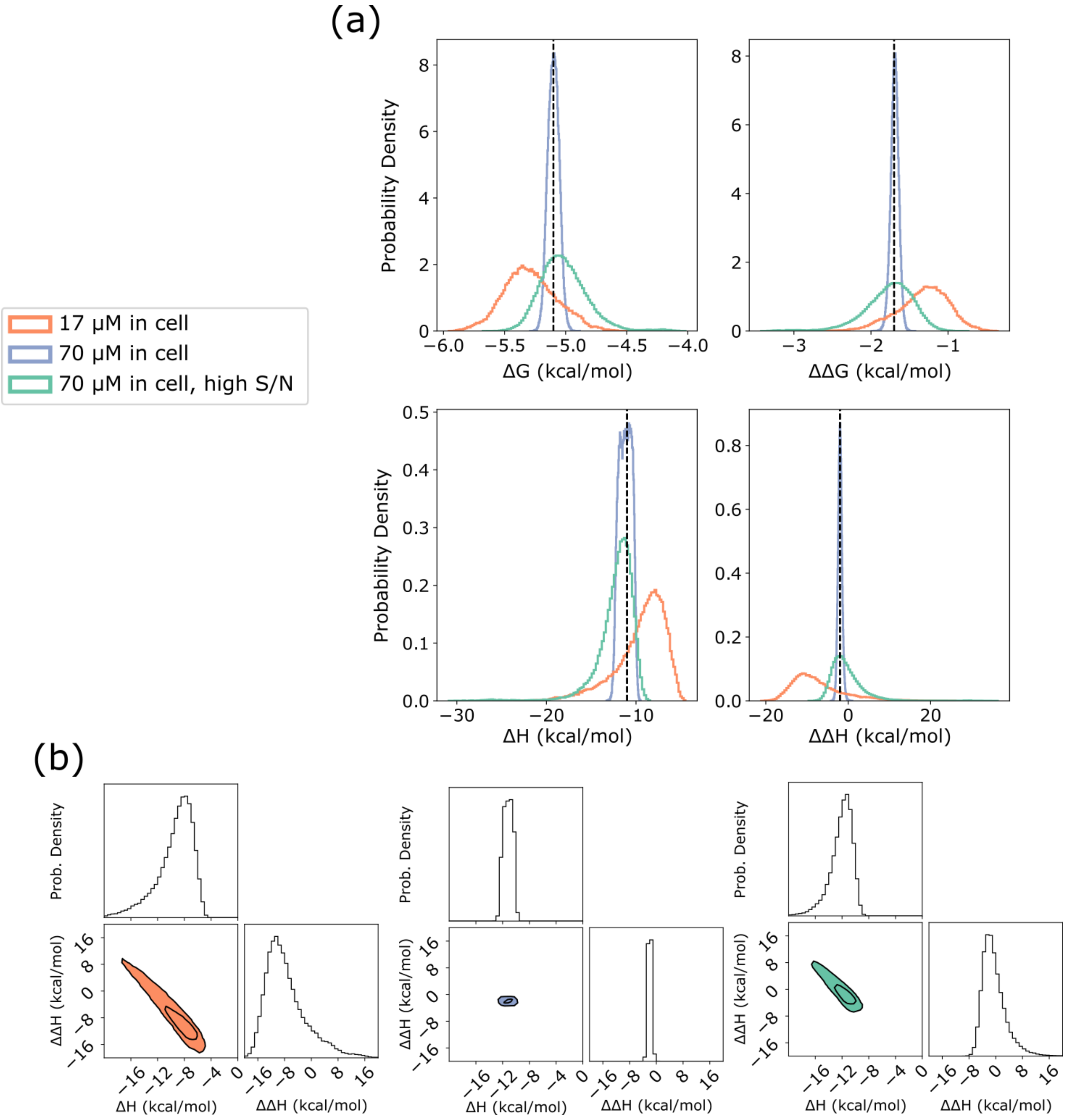


**S9 Figure: Marginal distributions for thermodynamic parameters for BSN-like synthetic isotherms.** (a) one-dimensional distributions for thermodynamic parameters for synthetic isotherms ($\Delta$G = -5.1, $\Delta\Delta$G = -1.7, $\Delta$H = -11, $\Delta\Delta$H = -2) generated with cell concentrations of 17 (orange) and 70 $\mu$M (blue and green). Syringe concentrations are 900 $\mu$M and 2000 $\mu$M respectively. The orange and blue isotherms are generated with noise taken from a gaussian distribution of width $\sigma$=0.2 $\mu$cal, while the green is generated with $\sigma$=0.8 ucal. (b) two-dimensional marginal distributions for the same models as (a), in the $\Delta$H-$\Delta\Delta$H dimension. Contours are drawn at 95 and 50% probability density. While raising the synthetic experimental concentration dramatically improves precision in all model parameters, much of this is due to the increased S/N ratio associated with the higher concentration. Scaling synthetic model noise with the increase in cell concentration reduces the precision of model enthalpies, although they are still narrower than the distributions for the low concentration isotherm.

| Isotherm | ∆G sum min | ∆G sum max | ∆H sum min | ∆H sum max | conc ratio min | conc ratio max |
|---|---|---|---|---|---|---|
| SPAG5 | -15.0 | -14.5 | -34 | -28 | 71 | 75 |
| BSN (I) | -12.2 | -11.7 | -29 | -21 | 64 | 86 |
| BSN (II) | -14.5 | -14.1 | -20 | -17 | 61 | 66 |
| SLC9A2 | -14.2 | -13.5 | -29 | -22 | 72 | 84 |
| Ebola VP35 | -15.7 | -15.3 | -27 | -22 | 21 | 22 |
| GLCCI | -13.2 | -12.6 | -21 | -16 | 85 | 99 |
| BIM | -18.2 | -16.1 | -26 | -22 | 25 | 28 |

**S1 Table: Credibility regions for 'sum' thermodynamic parameters and ratios of concentrations.** 95% credibility region from sampled posterior distributions for the ∆G sum(2∆G+∆∆G) and dH sum(2∆H+∆∆H) as well as the ratio of concentrations ([peptide]/[LC8]). Credibility regions for ∆G and ∆H sums are frequently narrower than the credibility regions for individual parameters (Table 1).

| Isotherm | ΔG min | ΔG max | ΔΔG min | ΔΔG max | ΔH min | ΔH max | ΔΔH min | ΔΔH max | -TΔS min | -TΔS max | -TΔΔS min | -TΔΔS max |
|---|---|---|---|---|---|---|---|---|---|---|---|---|
| SPAG5 | -6.93 | -6.2 | -2.12 | -1.13 | -19.11 | -12.59 | -1.87 | 3.7 | 5.97 | 12.61 | -5.78 | 0.69 |
| BSN (I) | -7 | -4.76 | -2 | 0.28 | -36.87 | -7.57 | -15.87 | 48.08 | 0.77 | 31.97 | -50 | 16.02 |
| BSN (II) | -7.09 | -6.44 | -1.22 | -0.39 | -6.41 | -4.44 | -9.75 | -6.53 | -2.22 | -0.46 | 5.68 | 8.99 |
| SLC9A2 | -6.85 | -5.47 | -2.64 | -0.53 | -24.97 | -9.75 | -5.19 | 24.76 | 3.19 | 19.49 | -27.42 | 4.6 |
| Ebola VP35 | -7.4 | -6.81 | -1.67 | -0.92 | -15.23 | -10.29 | -0.77 | 1.45 | 3.22 | 8.08 | -3.11 | -0.18 |
| GLCCI | -6.11 | -5.21 | -2.27 | -0.99 | -19.49 | -9.35 | -0.52 | 21.35 | 3.43 | 14.12 | -23.57 | -0.48 |
| BIM | -9.5 | -6.95 | -2.18 | 0.8 | -13.5 | -9.57 | -3.17 | -0.47 | 1.05 | 5.54 | -0.93 | 2.32 |

**S2 Table: Ranges for thermodynamic parameters for LC8-client binding when modeled with ±20% LC8 concentration.** Table 1 in the main text contains equivalent information at ±10% LC8 concentration. Values delineate 95% Bayesian credibility regions from sampled posterior distributions, when modeled with ±20% priors for LC8 concentration. Distributions are largely very similar to those presented in Table 1, with a slight decrease in precision. BSN I, for which posterior distributions are significantly broader, is the only notable exception.

| Peptide | ΔG | ΔH | *n* |
|---|---|---|---|
| SPAG5 | -8.0 ± 0.3 | -15.10 ± 0.09 | 1.01 ± 0.004 |
| BSN (I) | -7.3 ± 0.7 | -12.8 ± 0.3 | 0.99 ± 0.02 |
| BSN (II) | -7.9 ± 1.2 | -9.8 ± 0.3 | 0.95 ± 0.02 |
| SLC9A2 | -7.7 ± 0.6 | -11.52 ± 0.15 | 1.01 ± 0.01 |
| Ebola VP35 | -8.4 ± 0.4 | -11.84 ± 0.06 | 1.00 ± 0.004 |
| GLCCI | -7.3 ± 0.6 | -10.8 ± 0.2 | 1.00 ± 0.015 |
| BIM | -8.6 ± 0.7 | -11.45 ± 0.09 | 1.02 ± 0.006 |

**S3 Table: Binding parameters determined from identical sites model fits, as published in Jespersen et. al. (2019)(9).**

| Isotherm | # samples | # walkers | dG prior (kcal/mol) | dH prior (kcal/mol) | ddG prior (kcal/mol) | ddH Prior (kcal/mol) | X initial Prior (µcal) | M initial Prior (µcal) |
|---|---|---|---|---|---|---|---|---|
| Synthetic isotherm models | 50,000 | 25-50 | -3 to -10 | -50 to 0 | -4 to 4 | -40 to 40 | Varied - ±10% of stated unless otherwise noted | Varied - ±10% of stated unless otherwise noted |
| Single isotherm experimental models | 100,000 | 50 | -3 to -10 | -50 to 0 | -4 to 4 | -40 to 40 | ± 10% of stated | Up to ±50% of stated |
| Two isotherm experimental models | 200,000 | 50 | -3 to -10 | -50 to 0 | -4 to 4 | -40 to 40 | ± 10% of stated | Up to ±50% of stated |

**S4 Table: Model priors and sampling lengths for all isotherms.**

**Quantifying cooperative multisite binding in the hub protein LC8 through Bayesian inference, Supplementary Information**

**Best practices for the application of Bayesian statistical models to isothermal titration calorimetry**

**Aidan B Estelle[1], August George[2], Elisar J Barbar*[1], Daniel M Zuckerman*[2]**

[1]Department of Biochemistry and Biophysics, Oregon State University, Corvallis, Oregon 97331, United States

[2]Department of Biomedical Engineering, School of Medicine, Oregon Health and Science University, Portland, Oregon 97239, United states

This guide is intended to help anyone attempting to perform Bayesian inference on isothermal titration calorimetry (ITC) data for mono- or multivalent systems. The concentration degeneracy identified in the main paper makes the necessary Markov chain Monte Carlo (MCMC) even more challenging than usual. This guide is based on considerable effort experimenting with multiple algorithms and software packages.

*Sampling*

Adequate MCMC sampling is essential to the production of reliable parameter estimates from Bayesian inference. In our tests, we found the affine invariant sampling method (1) implemented in the python package EMCEE (2) performed by far the most effectively and efficiently for ITC modeling when including concentrations as model parameters. The affine method performs well in difficult parameter terrain (e.g. with a significant degeneracy), can run partially in parallel, and doesn't require the differentiation used in Hamilton Monte Carlo and related methods. The algorithm's employment of multiple MCMC walkers and construction of trial moves based on the locations of other walkers enables it to track narrow features in the landscape in an adaptive fashion.

EMCEE uses two 'hyperparameters' having to do with sampling: the number of walkers, and the number of steps each walker will take. While the EMCEE documentation recommends 3x the number of parameters as a minimum walker count, we found a higher number (e.g., 50 walkers for our 8 parameter model) improves the efficiency of sampling and reduces the likelihood of individual walkers "getting stuck," unable to move significantly relative to other walkers. For our model specifically, around 100,000 steps for each walker were sufficient for excellent sampling (confirmed by replicates), although this number is likely to be very model specific.

Although EMCEE's default trial move method, the 'stretch' move, performs robustly on ITC models, we found marginally increased performance through use of a mix at a ratio of 80:20% of the stretch move: the 'differential evolution snooker' move. The move performance may be model specific, however, so if possibly it is best to try a few trial move selection methods

Since there is no reliable way to predict how much sampling will be required, the quality of sampled posterior distributions must be checked. Methods for determining what qualifies as 'adequate' vary widely, such as measurement of r-hat (3), autocorrelation time (4) or effective sample size (5), but we suggest the use of replicate MCMC runs (each with its own seed) as a metric for sufficient sampling. When sampling is adequate, the individual replicate posterior distributions for each parameter – or any observable of interest – should converge to be near-identical to each other (see supp fig. 5 of this manuscript for an example).

The number of sample steps required also scales roughly with the number of model parameters, as each parameter geometrically grows the size of the parameter space that must be explored. This means that any increase in model complexity, or application of global models with additional nuisance parameters will increase the number of required sampling steps – both the number of walkers and steps per walker, in EMCEE.

*Global Models for multiple isotherms*

When applying a 'global' model to multiple isotherms (with the aim of inferring a single model governing all isotherms), the most significant consideration is the addition of new nuisance parameters. The heat of dilution ($\Delta H_0$ in our terminology), the model noise (A Gaussian distribution with mean = 0 and unknown variance sigma), and analyte concentrations are all isotherm specific parameters, so each isotherm will have its own set of these. In some cases, it may be possible to eliminate some parameters: e.g., If all isotherms are collected from the same stocks of protein, it could be acceptable to apply a 'global' prior for each concentration, rather than model each individual concentration for each isotherm. In general, though, a more complete model is better, and there is reduced risk of overfitting in a Bayesian framework relative to frequentist methodologies (6).

Experimentally, global models are best applied to isotherms with varied experimental conditions – particularly concentrations. While global models of technical-replicate isotherms are not useless, their ability to improve precision is limited, while isotherms containing non-overlapping information through variation of initial concentrations are more likely to improve global models (7, 8).

*Concentration Priors*

As discussed at length in the manuscript, individual concentrations cannot be determined through fitting or modeling. This means that limits set on the concentration ranges (i.e., by priors) are of particular significance to accurate modeling. Fortunately, the *ratio* of concentrations can be determined from the model (see main manuscript and Figs. 2 and 3), which functionally means that knowing only one concentration is adequate for a given experiment. Taking this into account, we advise that whichever experimental concentration can be determined with greater precision be used to limit the range of degenerate model solutions. The most naïve method to do this is to give the known concentration a uniform prior of ±X% around a measured value, where X is some measured uncertainty. Once that is set, the other analyte concentration can be set to as wide a prior as is needed: provided the two-dimensional posterior distribution shows a clear diagonal line within the parameter space, the model is able to find and sample along the degenerate line. If confidence in concentrations is particularly high, it may be reasonable to use gaussian distributions for priors, as it would likely be reasonable to expect Gaussian noise in concentration measurement.

*Experimental considerations*

While the normal considerations in setting up an ITC experiment apply, two additional considerations may apply for Bayesian modeling. First, as we discuss in the manuscript, when modeling complex systems, there is no longer a single affinity around which to design experiments with regards to the ITC parameter c ($c = n[\text{cell}]/K_d$). In such cases, we believe global isotherms collected at multiple concentrations are likely to contain all the information necessary to extract all thermodynamic parameters, although we did not systematically explore this design consideration.

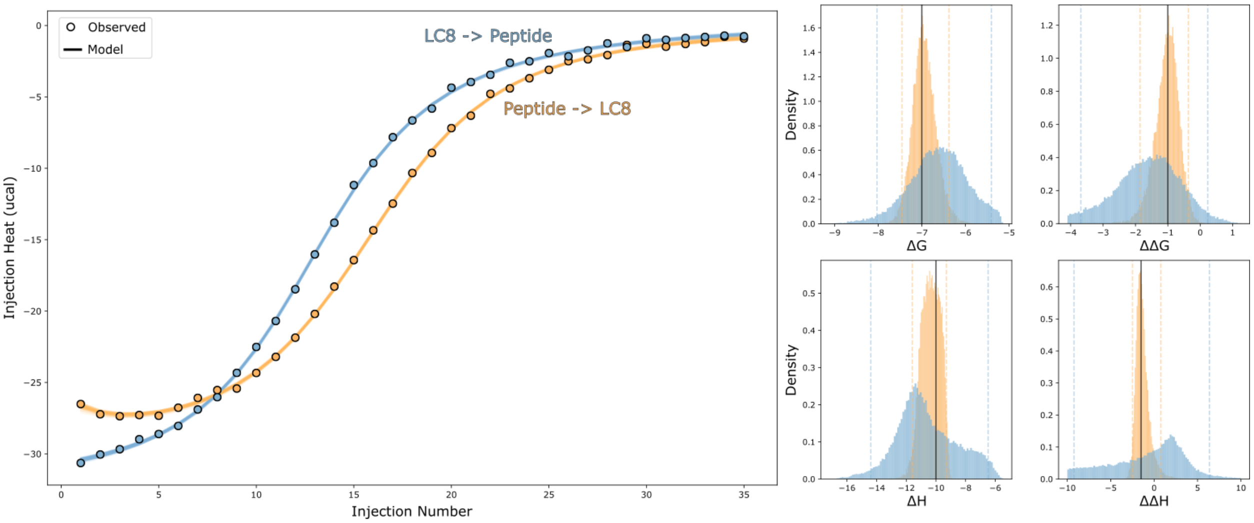


**Supplemental Figure S9: direction of injection impacts parameter determinability.** Simulated isotherms of injection of LC8 into peptide (blue), and peptide into LC8 (yellow). Posterior distributions for injection of peptide into LC8 are significantly narrower than those for the LC8 titration.

Second, the information content of isotherms is dependent on the direction of injection. In tests for our specific 2-site system, we found from synthetic data that injecting the ligand into a solution of the protein with two sites returns significantly narrower distributions (on the order of 2-3 kcal/mol) than the reverse. We believe this is likely to depend on the binding model being tested and is best tested with synthetic isotherms prior to experimentation. Our example notebook, available at https://github.com/ZuckermanLab/Bayesian_ITC includes code for generating and modeling synthetic isotherms.


1. J. Goodman, J. Weare, Ensemble samplers with affine invariance. *CAMCoS* **5**, 65–80 (2010).

2. D. Foreman-Mackey, D. W. Hogg, D. Lang, J. Goodman, emcee : The MCMC Hammer. *Publications of the Astronomical Society of the Pacific* **125**, 306–312 (2013).

3. A. Gelman, D. B. Rubin, Inference from Iterative Simulation Using Multiple Sequences. *Statistical Science* **7**, 457–472 (1992).

4. M. Wallerberger, Efficient estimation of autocorrelation spectra (2019) (June 10, 2022).

5. V. Elvira, L. Martino, C. P. Robert, Rethinking the Effective Sample Size. *Int Statistical Rev*, insr.12500 (2022).

6. A. Gelman, *et al.*, Bayesian Data Analysis Third edition. 677.

7. M. V. C. Cardoso, *et al.*, CALX-CBD1 Ca2+-Binding Cooperativity Studied by NMR Spectroscopy and ITC with Bayesian Statistics. *Biophysical Journal* **119**, 337–348 (2020).

8. H. Duvvuri, L. C. Wheeler, M. J. Harms, pytc: Open-Source Python Software for Global Analyses of Isothermal Titration Calorimetry Data. *Biochemistry* **57**, 2578–2583 (2018).
